# Critical structural perturbations of ribozyme active sites induced by 2’-O-methylation commonly used in structural studies

**DOI:** 10.1101/2025.05.26.656154

**Authors:** Sélène Forget, Guillaume Stirnemann

## Abstract

Most naturally occurring ribozymes catalyze self-splicing reactions through a 2’-OH group. Consequently, experimental structures of precatalytic states often require chemical modifications of the 2’-OH, such as its removal or methylation. However, the impact of these chemical modifications on the active site structure remains largely unexplored, which raises important questions since methylated structures are often taken as being representative of pre-catalytic states. Here, we employ extensive atomistic simulations critically compared to and fine-tuned on experimental data, and we revisit experimental results to show that 2’-O-methylation critically affects reactant geometries and, therefore, the possible reaction mechanisms inferred from the structures. Our results also challenge the common assumption that 2’-O-methylation stabilizes the C3’-endo puckering conformation. Our findings, consistent with recent experimental data on ribosome structure, reveal that this effect is highly sensitive to the local secondary structure and is often overstated. For three investigated small-cleaving ribozymes, the C2’-endo conformation observed for chemically-modified active site residues through 2’-O-methylation is not stable upon methyl group removal to obtain the catalytically-relevant hydroxylated state. This suggests that these geometries arise primarily from a combinaison of steric hindrances and electrostatic interactions with the surrounding environment rather than intrinsic conformational preferences of the ribose upon methylation.

## 1 Introduction

The ability to catalyze chemical reactions is one of the remarkable features of certain ribonucleic acids, known as ribozymes ^1,2^. However, the range of chemical reactions catalyzed by ribozymes remains significantly narrower compared to protein enzymes ^3–6^. Most known ribozymes primarily catalyze reactions such as self-ligation or selfsplicing. These reactions play a crucial role in viral replication and RNA splicing mechanisms ^3,7,8^.

Self-splicing is a substitution reaction at a phosphate group, where a conventional 3’-5’ phosphodiester bond is replaced by a 2’-3’ cyclic phosphate linkage. This process begins with a nucleophilic attack by the 2’-hydroxyl group of the same nucleotide ^9^, and is facilitated by proton transfers and active-site conformations that promote the desired reaction while suppressing side reactions.

**Figure 1:**
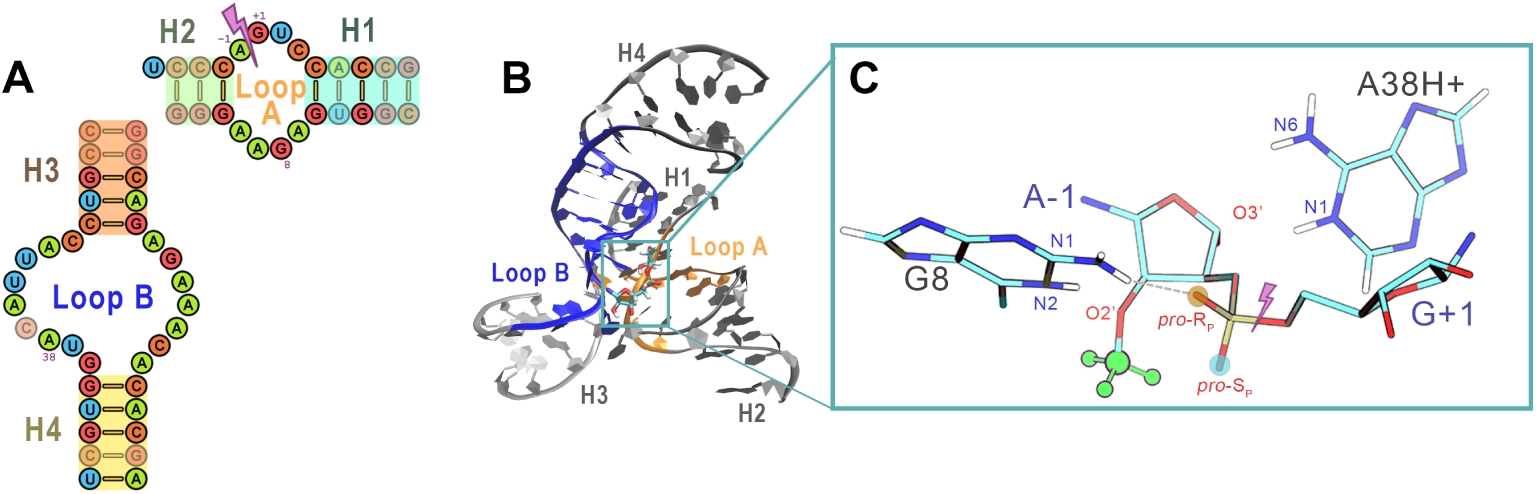
Overview of the minimal hairpin ribozyme (HpR) analyzed in this study. (A) Sequence and secondary structure showing helices H1-H4, loops A and B, and the cleavage site (pink arrow in loop A). (B) Crystal structure (PDB 2oue) using the same labels. (C) Zoom on the active site structure of the O2’-methylated crystal structure, with added hydrogens as described in the Methods section.

To experimentally capture ribozymes in their precatalytic state, the reaction must be inhibited—typically through chemical modification ^10,11^. Among such systems, the hairpin ribozyme (HpR) is one of the most thoroughly studied using this approach ^11–15^. For this ribozyme, common inhibitory strategies include reducing or eliminating the reactivity of the 2’-OH group via dehydroxylation ^11,16,17^, methylation ^11,12,17–20^, or mutating key catalytic residues that are directly or indirectly involved in the reaction mechanism ^11,18,21^.

This methodology parallels strategies used in enzyme studies, where substrate analogs are chemically altered to inhibit activity. However, there are two important differences: first, for protein enzymes, it is typically the substrate, and not the protein scaffold, that is chemically altered. And unlike proteins, whose conformational landscapes are often considered more rigid, ribozymes may be more susceptible to structural changes from such modifications because they typically evolve along more flexible conformational landscapes. A fundamental paradox arises: while structural studies frequently assume that 2’-O-methylation does not significantly alter active site geometry, this same modification is a widespread and biologically important RNA post-transcriptional modification (PTM) ^22–27^, known to regulate numerous cellular processes ^25,28–39^. Mounting evidence suggests that even single methylation events can trigger substantial structural rearrangements, either locally or across broader regions of the RNA ^40–46^. Yet, beyond a few well-characterized examples, the full extent of these structural consequences remains poorly understood.

In organic chemistry, it is well established that the conformation of cyclic structures is highly sensitive to the nature of their carbon substituents. For nucleic acids, the three-dimensional conformation of the ribose ring in each nucleotide is called the sugar puckering ^47,48^. This puckering describes how individual atoms within the five-membered sugar ring deviate from planarity, depending on which carbon atoms are displaced above or below the ring’s plane. Ribose puckering is highly sensitive to the chemical environment, with conformational preferences influenced by steric effects and intra- or intermolecular interactions.

The dominant view in the literature is that 2’-O-methylation modulates ribose puckering ^49^, typically shifting the equilibrium toward the C3’-endo conformation ^40,50–54^, which is the most common sugar pucker in RNA duplexes ^55–58^. This methylation-induced bias contributes to duplex stabilization and can drive unpaired regions to adopt helical conformations ^40,59^. However, exceptions to this pattern are increasingly recognized. For example, in several crystal structures of precatalytic RNA, 2’-O-methylated riboses have been observed to adopt the C2’-endo conformation ^12,60–63^, highlighting the complexity and variability of RNA conformational behavior.

In addition, a closer examination of the existing literature, particularly on the HpR, reveals a more nuanced and, at times, conflicting picture.

First, molecular dynamics (MD) simulations initiated from the canonical crystal structure of the methylated precatalytic state consistently revealed a transition from the C2’-endo to the C3’-endo conformation upon removal of the methyl group, across multiple force fields ^64–68^.

Second, while most precatalytic HpR structures exhibit both 2’-O-methylation and C2’-endo puckering, structures featuring chemical modifications at other positions display a variety of puckering states, including C3’-endo ^11,16,18,21^.

Third, recent cryo-EM structures of the human ribosome, which includes numerous 2’-O-methylated nucleotides, have shown that riboses in unpaired regions frequently do not adopt the C3’-endo conformation ^69–71^.

Finally, the tendency of isolated nucleotides to favor the C3’-endo conformation due to methylation appears overstated in the literature. Experimental NMR measurements indicate that the stabilizing effect of methylation is relatively modest (no more than 0.1 kcal/mol) corresponding to a population shift of only 5–10% ^50^. This is far less pronounced than sometimes implied.

These observations prompted us to ask whether 2’-O-methylation universally promotes the C3’-endo conformation relative to unmodified ribose. Focusing on the active site of the HpR, we use molecular dynamics simulations to demonstrate that, in this specific context, 2’-O-methylation exerts the opposite effect, challenging the prevailing view in the literature. Our findings remain robust across multiple force fields, particularly when simulations are refined using available experimental data, and similar behavior is observed in other ribozymes. This context sensitivity of the puckering sensitivity to 2’-O-methylation reconciles previously conflicting experimental results and has important implications for the interpretation of chemically modified crystal structures and the mechanistic understanding of ribozyme catalysis.

## 2 Materials and methods

### 2.1 Simulation and analyses: generalities

All simulations, from the system preparation to production runs, were performed with GROMACS 2022.3^72^, patched with PLUMED 2.7^73,74^. Input files required to reproduce these simulations will be made available on the online Zenodo repository. The home-made scripts developed for the simulation analysis can be found online on the GitHub repository of the group, as referenced at the end of the article. These codes require both the Python open-source MDAnalysis ^75,76^ and Barnaba ^77^ libraries. Visualization and atomistic views were done with the VMD software ^78^, combined with the Collective Variables Dashboard ^79^. The converged part of the simulations, which were produced using replica exchange methodologies, was analyzed using YACARE ^80^. This clustering tool was set up to identify clusters of frames that represented more than 2% of the total. The clustering was performed over the data space of distances between each pair of nucleobases, given that these distances could be considered as contacts formed during a significant fraction of the time (i.e., under 5 Å in at least 5% but less than 95% of the analyzed frames). This allows to select structural features that can distinguish between different conformations.

### 2.2 Forcefields

Two forcefields were used in this paper : Amber14_ff99bsc0 *χ*_OL3_*eζ*_OL1_ ^81^ (with TIP3P water), which derives from the AMBER ff99 force field ^82^, and was subsequently corrected ^81,83,84^, and DES-Amber 3.20^85^ with TIP4P water. These two forcefields were selected to represent two important families of nucleic acid force fields, while having shown to be reasonably adapted to simulations of the HpR system ^67,68^.

These forcefields lacked certain non-standard residues which were required for the simulation set up, notably the protonated adenine (RAP) residue. The corresponding parameters were adapted from the work of Mlynský et al. ^65^. The complete topology used in these simulations is provided in our previous work ^68^. Methylated adenosine was adapted from the parameters provided by Mlynksy et al. ^65^ in order to incorporate this residue in Amber14_ff99bsc0*χ*_OL3_*eζ*_OL1_. For both residues, DES-Amber parameters were obtained by deriving the standard equivalent residue (adenosine) on which the charge redistribution was obtained by the same charge correction as in the Amber force field. The final residues parameters can be found in the Supporting Information.

### 2.3 Ribozyme simulations

Solution simulations were performed using the same protocol as described in our previous work ^68^, starting from a precatalytic state structure of the minimal hairpin (HpR) ribozyme (PDB 2oue) ^11^, the hammerhead (HhR, PDB 3zp8) ^86^, or the apo structure of glmS (glmS, PDB 3g8s) ^62^. For the HpR, following previous studies ^67,87^, A38 was protonated at the N1 position and G8 was left neutral. Other bases were taken in their standard protonation state at neutral pH. Crystallographic anions (SO^2^*^−^* and [Co(NH_3_)_6_)]) were removed; crystallographic water molecules were preserved, and the system solvated in a 0.2-M KCl solution. For HhR and glmS, all bases were considered in their standard protonation state at neutral pH, all crystallographic anions removed, the water molecules conserved, and the system was neutralized with sodium ions (which corresponds to the conditions of crystallization) and then solvated with water. Both systems were simulated without magnesium ions. All solvated ribozymes were then subject to minimization, equilibration, and production runs in the NPT ensemble at 300 K and 1 bar using the workflow described before ^68^.

Briefly, the minimization steps were conducted in three phases, withn first positional restraints of 1,000 kJ/mol/nm² imposed on all backbone atoms. These steps include 5,000 energy minimization iterations, followed by two 500 ps equilibration phases under NVT and NPT conditions, respectively, at a temperature of 300 K and pressure of 1 bar. Thermal and pressure control was achieved using Parrinello-Rahman barostat and the velocity-rescaling thermostat ^88^l. To gradually accommodate the structural relaxation of the nucleic backbone, restraint forces are systematically reduced across three sequential steps, decreasing from 1,000 to 100 to 10 kJ/mol/nm². After the restraints are lifted, the system undergoes unrestrained simulations: first under NVT for 100 ps, followed by 100 ps under NPT conditions. A comprehensive description of the equilibration strategy and the associated parameter files for each stage will be made available in a Zenodo repository.

Electrostatic interactions are evaluated using the particle-mesh Ewald (PME) approach ^89^, while Lennard-Jones forces are calculated within a 10.0 Å cutoff. The LINCS algorithm ^90^ is utilized to constrain all bonds involving hydrogen atoms. A simulation timestep of 2 fs is consistently used throughout. The PME grid is constructed with a spacing of 0.12 nm, employing cubic interpolation for charge distribution.

For HpR and HhR, enhanced sampling simulations were performed using a Hamiltonian replica exchange scheme, here Replica Exchange with Solute Tempering (REST2) ^91,92^. We used 24 replicas with rescaling factors *λ* ranging from 1 to 0.667, attempting exchanges every 1 ps, at 300 K, for simulation lengths varying from 500 ns to 1 *µ*s. Convergence was evaluated through block averaging of catalytic site conformation populations and monitoring the evolution of the eRMSD metric ^93^ over time for the unscaled replica (see Figure S1 and S2 for the respective time convergences). The average exchange rate was close to 15-22%.

For crystal simulations (performed with Amber14_ff99bsc0 *χ*_OL3_*eζ*_OL1_), we chose to adopt a naive approach and reduce the complexity of the system to a minimum, aware that many choices can have a complex and significant impact on the results ^94,95^. We did not introduce any ions or additional compounds present in the medium during the crystallization process, but instead added TIP3P water molecules with Na^+^ counter ions to neutralize the system. We precise here that among these chemicals (poly(ethylene glycol) methyl ether, spermidine, sodium cacodylate, LiSO_4_, [Co(NH_3_)_6_)], only crystal water molecules were kept. The rest of the water was added using Gromacs’ `solvate` command. We chose to simulate the crystal under the conditions of its growth, i.e., at 300 K and 1 bar.

The crystal lattice was reconstructed according to space group P6(1)22, where twelve symmetry-related copies of the ribozyme make up one unit cell. We constructed a supercell using Chimera 1.17.3 software ^96^, yielding a triclinic box of size 93.272 × 93.272 × 131.250 Å and angles 90 × 90 × 120°. We verified that the cell volume remained stable during the NPT simulations, with a standard deviation between ∼ 0.013% and ∼0.017% during the entire trajectory.

The rescaled “hot region” consists of only one of the twelve ribozymes of the lattice, that is the only one analyzed (see the elements displayed in licorice full-atom style using VMD ^78^ representation in Figure S3 B, C.) The 32 replicas in our setup spanned rescaling factors *λ* ranging from 1 to 0.667, achieving an exchange rate of 22%, and the simulation ran for 760 ns. Minimization, equilibration and production steps were conducted similarly to the bulk simulations, as described above.

### 2.4 Mono and di-nucleotides

To properly assess the reliability of our force fields, we tried to reproduce the experiments conducted by Kawai et al. ^50^. We simulated isolated uracil mononucleotides and dinucleotides in the bulk, in conditions similar to the experiments. We also adopted their nomenclature: U stands for the uridine nucleoside, Um for 2’-O-methylated uridine. The dimers are composed of two nucleotides joined by a phosphodiester bond, which is why the letter “p” is introduced in between the residue names. For instance, UmpU refers to two uracils with the 5’ residue methylated on its 2’-O atom.The force field topology files already included parameters for isolated nucleosides (U, residue type RUN) and 3’ and 5’ termini nucleotides (UpU, residue types RU5 + RU3). It should be noted that the methylated nucleotides are not standard residues and are not provided by the original force field. The new residue types were created using the same methodology as outlined above for the methylated adenosine and protonated adenosine. The final charge distributions can be found in the Supporting Information. In order to set up Um, we created a new residue type MUN, which constructed from the uracil nucleoside RUN to which we added the methyl group. To set up UmpU, we created a 5’ termini methylated uracil and the corresponding new residue type MU5. The 3’ hydrogen atom was eliminated from MUN, and the charge discrepancy between the 5’ uracil termini (RU5) and the uracil nucleoside was employed to generate the 5’ methylated uracil termini, MU5.

The mononucleotides and dinucleotides were solvated in a cubic box of 3 nm of TIP3P water molecules at 300 K, together with potassium counter ions to neutralize the system. The minimization and equilibration steps were conducted as described above for the ribozymes, with the Amber14_ff99bsc0 *χ*_OL3_*eζ*_OL1_ forcefield.

OPES metadynamics simulations ^97^ were performed to sample the puckering pseudo-rotation angle equilibrium. Howerver, due to software constraints, we did not apply a bias along the puckering angle but instead targeted the *τ*_2_ dihedral angle, following the methodology of Mlynsky et al. ^67^. To validate that *τ*_2_ is a good proxy for the puckering angle, we examined the correlation between the dihedral angle with the puckering pseudo-rotation angle in the REST2 bulk simulations of the methylated HpR (Figure S4). Indeed, negative *τ*_2_ values correspond to C2’-endo puckering conformations, while positive *τ*_2_ values align with C3’-endo puckering conformations. Since free-energy differences between the two conformations are obtained by integration of the populations in each of these basins, the system thermodynamics is not affected.

For each system, we launched 250 ns of OPES simulation for each of the mononucleotides and dinucleotides, and set the BARRIER parameter at 20 kcal/mol. OPES simulation convergence was assessed by comparing profiles in which early portions of the trajectory of different sizes were removed. We calculated the standard deviation for the ratio *p_C_*_3_*t_−endo_*/*p_C_*_2_*t_−endo_* Each OPES trajectory was truncated by its first *x* percent and we plotted this ratio as a function of *x* (Figure S5). These graphs reveal an approximate sweet spot, where the sd is minimal, typically for *x*=20%. This corresponds to the region where the potential has reached a quasi-static behavior, and the free energy profile is converged. Based on these observations, we systematically removed the first 20% of the trajectories and considered a reasonable sd to be below 0.2 kcal/mol for the ΔGs.

### 2.5 Local bias of the pseudorotation angle for 2’-O-methylation

The bias on the *τ*_2_ dihedral angle designed as a local correction of the HpR methylated ribose was defined as an external potential defined by a free energy grid which covers the entire range of the *τ*_2_ dihedral angles. The grid was built from a target free energy profile which verified the 60/40 ratio experimentally observed in the NMR of the mono-nucleotide uracil ^50^.

This bas was built using a polynomial function of the form:

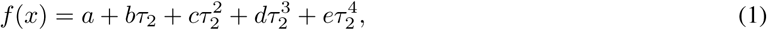

The *a*, *b*, *c*, *d* and *e* coefficients were determined by fitting the polynomial to the pass through theoretical points which met our criteria. One point was located at the *τ*_2_ value for the C3’-endo minimum, and fixed at 0 kcal/mol. Another was located at the *τ*_2_ value of the C2’-endo minimum. The other fitting points were extracted from the unmodified methylated mono-uracil OPES simulation: at the extremities of the explored range, and around the maximum of the free energy barrier. The points were adjusted empirically to obtain the desired free energy barrier and the 60/40 ratio, that is to say a difference of 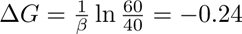 kcal/mol between the two local mini of the curve and a maximum identical to the original profile. The final grid bias is shown in red in Figure 3.

### 2.6 Creation of the additional dihedral terms for systematic corrections of the RNA forcefield

The dihedral potential parameters *K*, *n* and *δ* for these additional terms were determined from the shape of the grid bias described previously, and notably by the position of the minima and maxima of the bias curve. We represented in Figure S6 the grid bias curve (in black dots) together with the potential energy curve (in blue) that results from the terms we added to the force field. Because of the RNA sugar topology, the *τ*_2_ dihedral angle are never out of the [-60,60]° range. This facilitated the construction of the additional terms as the shape of the profile outside that range did not matter.

## 3 Results

### 3.1 Effect of methylation of the A-1 puckering angle

We begin by examining the puckering pseudo-rotation angle of the A-1 ribose in the minimal HpR in its precatalytic state, comparing its behavior when the ribose is methylated (as observed in most crystal structures ^11,12,17–20^) versus when it is in a regular hydroxylated state. Our computational approach is described in detail in the Methods section. Briefly, we performed atomistic molecular dynamics simulations in explicit solvent, employing two different forcefield combinations, Amber14_ff99bsc0*χ*_OL3_*eζ*_OL1_ ^81,82,84^ with TIP3P water and DES-Amber 3.20^85^ with TIP4P water.

Ideally, the conformational landscape of the A-1 ribose puckering could be mapped using enhanced sampling simulations along this collective variable ^98^. However, we found that such simulations did not converge even over long timescales. This suggests that puckering is not an intrinsically slow coordinate in this system but likely responds to changes in other, as-yet-unknown slow variables.

To facilitate conformational space exploration without relying on a specific choice of collective variable (CV), we performed Replica Exchange simulations with Solute Tempering (REST2) ^92^. This approach has been extensively applied in our previous studies on biomolecular systems ^99–103^, including RNA (by us ^104–106^ and others ^107–109)^ and this ribozyme ^68^. In the case of the HpR, we have demonstrated that while standard brute-force MD simulations are highly sensitive to the initial configuration of the unmethylated, precatalytic ribozyme (derived either from the methylated crystal structure or chemically guided constrained simulations), REST2 simulations achieve convergence on the sub-microsecond timescale regardless of the starting conformation ^68^. This allows for reliable estimation of the A-1 ribose puckering state.

Starting from the typical methylated crystal structure, with canonical protonation states for active-site residues, we simulated the A-1 2’-OH versions of the ribozyme (Figure S1). Rapid repuckering of the A-1 ribose from the C2’-endo conformation (observed in the crystal structure) to the C3’-endo conformation was observed with both forcefield combinations, consistent with prior simulation work using a broad spectrum of forcefields ^66–68,110^. However, transient repopulation of the C2’-endo state occurred during the simulations. As noted before, we are not aware of any forcefield that would lead to an opposite trend, and we previously showed that sampling long-enough or using enhanced sampling methods systematically resulted in a population of C3’-endo conformations close to 100% for the unmethylated structure ^68^.

From these simulations, we estimated the free-energy differences between the C2’-endo and C3’-endo conformational basins (see Methods). Using the Amber14_ff99bsc0*χ*_OL3_*eζ*_OL1_ forcefield, we observed a free-energy difference of approximately 3.9 kcal/mol favoring the C3’-endo conformation, while the DES-Amber 3.20 forcefield yielded a smaller difference of approximately 2.3 kcal/mol (Table 1).

**Table 1:** *p_C_*_3_*t_−endo_/p_C_*_2_*t_−endo_* ratios and the corresponding Δ*G* values calculated from the *τ*_2_ dihedral distribution in the converged portions of the REST2 simulations of the HpR, comparing different variants and conditions. Error bars correspond to standard deviations obtained from block-averaging.

| System | Parametrization | C3'-endo/C2'-endo | $\Delta G$ (kcal/mol) |
| --- | --- | --- | --- |
| Solution demethylated PC | $\chi_{OL3}\epsilon\zeta_{OL1}$ | 1074<sup>1</sup> | -3.9<sup>1</sup> |
| Solution demethylated PC | DES-Amber 3.20 | 58 ± 36 | -2.3 ± 0.3 |
| Solution methylated PC | $\chi_{OL3}\epsilon\zeta_{OL1}$ | 8.2 ± 3.6 | -1.2 ± 0.3 |
| Solution methylated PC | DES-Amber 3.20 | 1.1 ± 0.1 | -0.1 ± 0.05 |
| Crystal methylated PC | $\chi_{OL3}\epsilon\zeta_{OL1}$ | 18 ± 9 | -1.6 ± 0.3 |

<sup>1</sup> Error bars for this run could not be obtained as in some part of the trajectory, the C2'-endo conformation was not observed, resulting in infinite ratios.

When extending the solution simulations to the methylated state corresponding directly to the crystal structure, sharp differences emerge (Figure S1). First, while C3’-endo conformations begin to be explored, this occurs over much longer timescales compared to the unmethylated case, suggesting that the barrier for transitioning from C2’-endo to C3’-endo is significantly higher. After an equilibration period, the populations in each state stabilize, allowing us to estimate the free-energy differences between the C3’-endo and C2’-endo conformations. For the methylated ribose, the Amber14_ff99bsc0*χ*_OL3_*eζ*_OL1_ forcefield yields a free-energy difference of approximately 1.2 kcal/mol favoring C3’-endo, whereas the DES-Amber 3.20 forcefield produces a near-equilibrium scenario with a difference of only ≈ 0.1 kcal/mol, with roughly equal populations of the two conformations (Table 1).

These results indicate a 2–2.5 kcal/mol shift in favor of C2’-endo when transitioning from hydroxylated to methylated A-1 2’-OH ribose, with quantitatively similar trends among the two forcefields. With the DES-Amber 3.20 forcefield, this leads to an equal distribution of C2’-endo and C3’-endo conformations for the methylated ribose, whereas the C2’-endo conformation is almost negligible for the hydroxylated state. This underscores the profound impact of methylation on the conformational landscape of the HpR active site in solution.

### 3.2 Effect of crystallization of the A-1 puckering angle

We next consider whether changes in the A-1 puckering angle might arise due to the crystalline nature of the experimental structure. To investigate this, we directly simulated the methylated HpR in its crystal state, employing a carefully designed simulation strategy to mimic the crystal environment (see Methods, Figure S3). Interactions involving one ribozyme within the crystal lattice were perturbed in the REST2 scheme, with all subsequent analyses focused exclusively on this particular individual. Simulations were conducted under the same thermodynamic conditions as the solution simulations for consistency.

As expected, the crystal environment proved to be far more constrained than aqueous solution. Deviations from the reference crystal structure were significantly reduced, with large-amplitude motions of the H2 and H4 helix termini being restricted due to crystalline contacts (Figure S7). Despite these constraints, the A-1 puckering angle distribution remained remarkably consistent with that observed in solution (Figure S1 and Table 1).

It is worth noting that crystal simulations can be highly sensitive to the chemical composition of the system, which is often difficult to fully replicate and simulate ^94,95^. While alternative simulation strategies might yield differing results, it is striking that, despite large-scale differences in flexibility (Figure S1), the local structure of the active site appears unaffected by the crystalline environment (Table 1).

Together with our comparisons of the methylated and hydroxylated ribozyme variants in solution, we conclude that the differences between the methylated crystal structures and solution simulations of the unmethylated ribozyme are most likely due to the direct effects of methylation, rather than artifacts of the crystal environment.

### 3.3 Comparison with experimental data

The comparison with experimental results provides further support to the conclusion that methylation of the reactive ribose may in that context favor C2’-endo conformation over the C3’-endo. First, we observed that available PDB structures of the HpR in its pre-catalytic state show a marked preference for C2’-endo conformations when the reactive ribose is methylated (see Figure 2). It is noteworthy that while the majority of precatalytic minimal hairpin structures are derived from inactivation strategies involving methylation of the A-1 2’-OH ^11,12,17–20^ (and occasionally mutations at other residues), a few structures have been obtained using alternative approaches. In some cases, the 2’-OH was dehydroxylated ^11,16,17^, yielding a DNA analog of A-1, while in others, mutations were introduced in residues within the active site, excluding the A-1 ribose ^11,18,21^. Strikinly, the preferential adoption of the C2’-endo conformation is clearly associated with either 2’-OH methylation or dehydroxylation (Figure 2). In contrast, when the A-1 ribose remains chemically unaltered, the ribose conformation does not distinctly favor either the C2’-endo or C3’-endo basin or resides predominantly within the C3’-endo basin (Figure 2).

**Figure 2:**
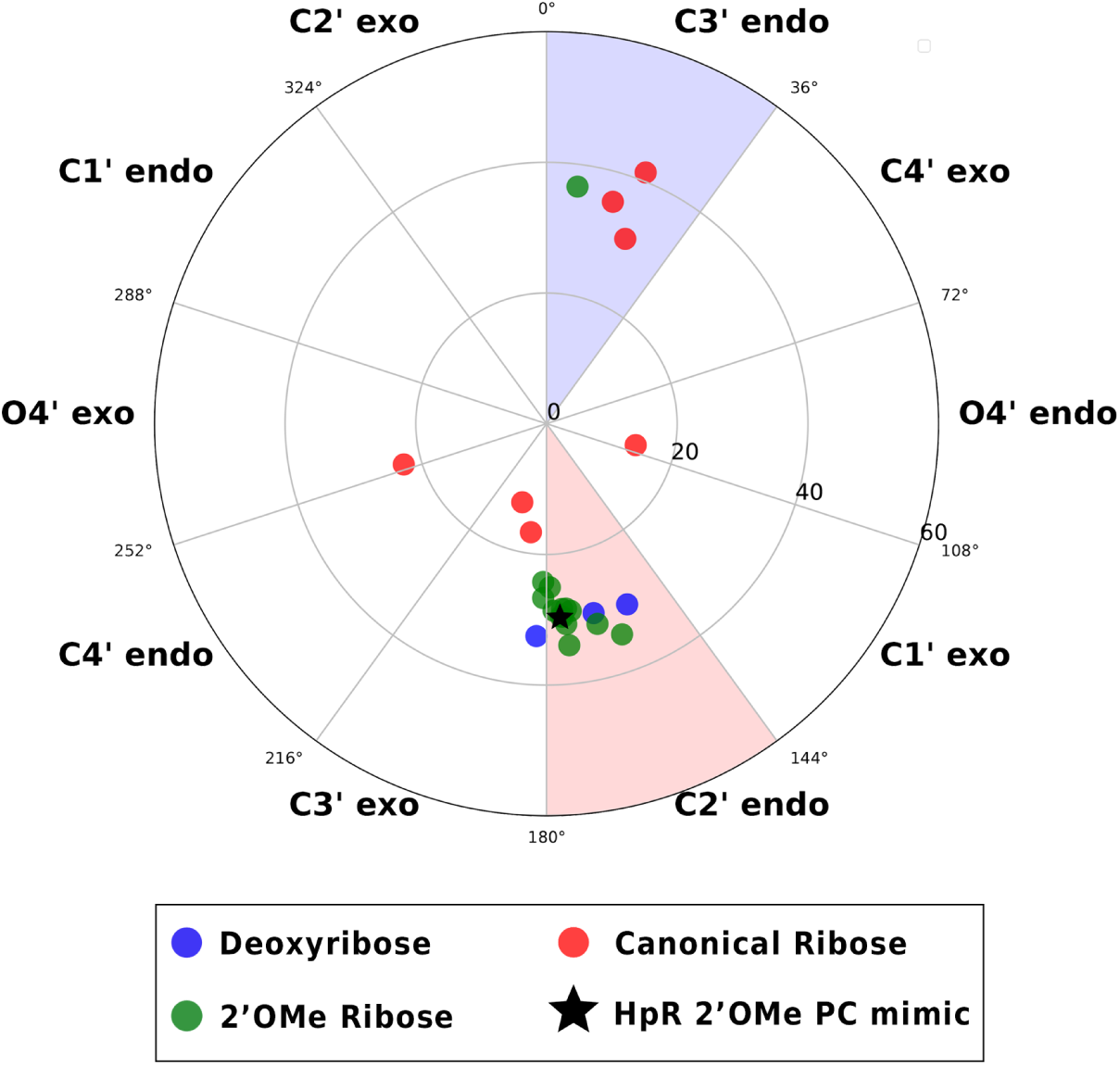
Overview of the puckering conformations of all the ligated crystal structures of HpR available in the PDB, colored according to the nature of the chemical modifications of their respective A-1 residues. The inhibition strategies used to block self-cleavage and mimic the pre-catalytic state are either methylation of the O2’ atom (A-1 2’OMe, in green) ^11,12,17–20^, removal of the O2’ atom (A-1:O2’ deoxy, in blue) ^11,16,17^, or chemical modification of another catalytic residue and A-1 kept unmodifed (in red) ^11,18,21^.

Second, we assess whether the forcefields used in our simulations accurately capture the puckering angle landscape, both with and without 2’-OH methylation. To the best of our knowledge, the only relevant experimental data come from NMR measurements of pyrimidine mono- and dinucleotides ^50–52^. Both RNA and methylated RNA slightly favor the C3’-endo conformation over the C2’-endo form, with methylation introducing only a minor additional stabilization, approximately 0.1-0.2 kcal/mol. Notably, larger effects reported in the literature often reflect enthalpic contributions alone rather than total free energy differences ^40,50^, as we discuss later in this manuscript.

To evaluate our forcefields, we simulated monoand di-uracyl systems ^50^ and computed the free-energy landscape along the *τ*_2_ dihedral angle of A-1, which directly correlates with the puckering pseudorotation angle, using the OPES-metadynamics enhanced sampling method (Figure S4). Although discrepancies between simulation and experiment never exceed 0.6 kcal/mol (Table 2), two key observations emerge: (i) neither regular nor methylated RNA fully reproduces the experimental balance between C2’-endo and C3’-endo conformations, and (ii) across all systems and forcefields, methylation consistently destabilizes the C3’-endo conformation relative to the unmethylated case, opposite to experimental findings.

**Table 2:**
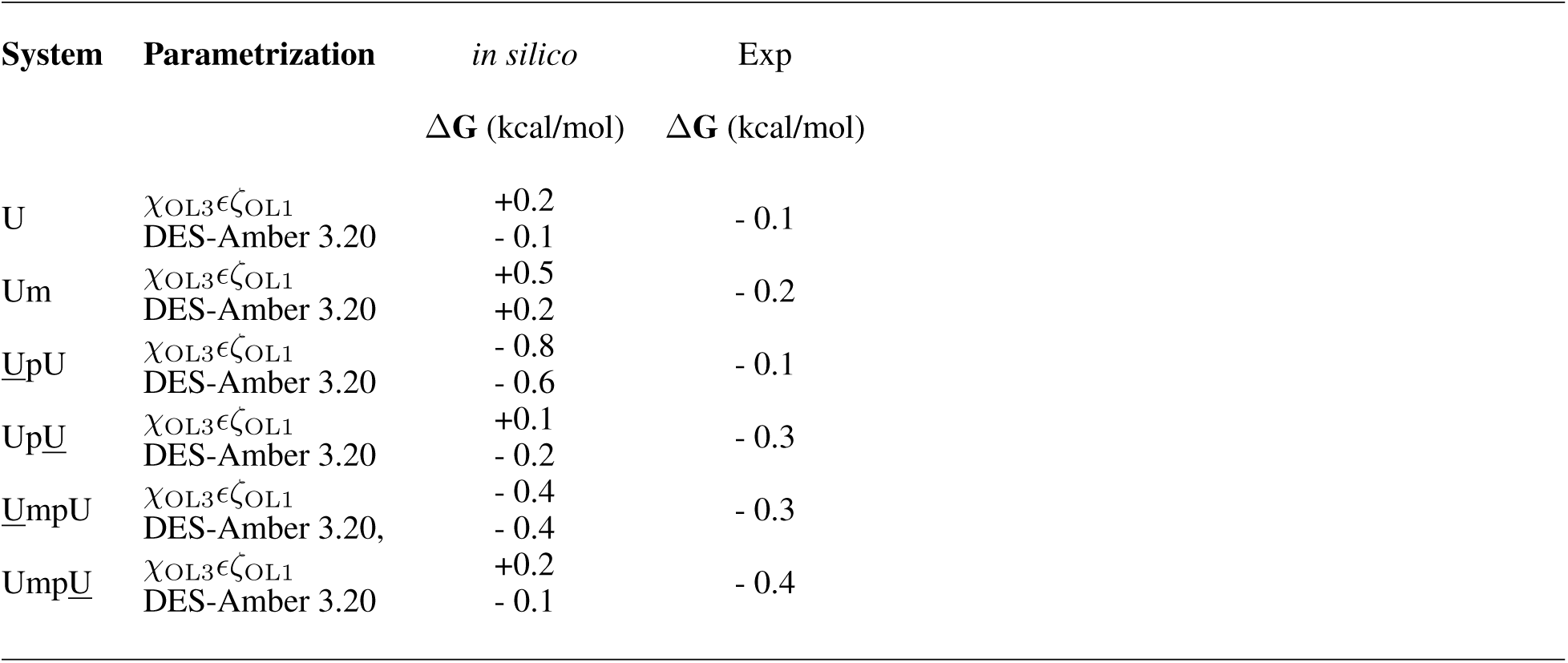
C3’-endo/C2’-endo puckering conformations equilibrium with the corresponding ΔG free energy differences in uridine mononucleosides, nucleotides and dinucleotides. The NMR experimental values are taken from ref ^50^ and *in silico* values are calculated from the *τ*_2_ dihedral distribution of our OPES simulations, excluding the first portion of the trajectories (Figure S5). For dinucleotides, we underline the considered ribose.

Among the tested forcefields, DES-Amber 3.20 performs best, showing the closest agreement with experimental data. It reproduces the C3’-endo/C2’-endo ratio within 0.1–0.3 kcal/mol, outperforming in that respect Amber14_ff99bsc0*χ*_OL3_ *eζ*_OL1_.

These simulations also underscore that while the intrinsic puckering preferences of isolated ribose influence conformational bias in the ribozyme active site, the overall free-energy landscape is primarily shaped by interactions with the surrounding environment. This is evident from the significantly different C3’-endo/C2’-endo free-energy differences observed in the ribozyme (Table 1) compared to isolated nucleotides (Table 2), and from the much larger forcefield-dependent variations in the ribozyme context.

### 3.4 Reparametrization of the methylated A-1 pseudo-rotation angle

In light of the observations above, it is however evident that the forcefields used do not accurately capture the effect of methylation on the puckering angle. Specifically, the stabilization of the C2’-endo conformation observed in our ribozyme simulations may stem from an inaccurate description of methylation, which experimentally is known to slightly favor the C3’-endo conformation in isolated pyrimidine nucleotides relative to their unmethylated RNA counterparts.

Although it is unlikely that the 2–2.5 kcal/mol shift upon 2’-O-methylation observed in our simulations can be fully explained by a 0.4–0.5 kcal/mol discrepancy in the methylation effect, we attempted to reparameterize one of the forcefields to better reproduce experimental puckering preferences. We selected the Amber14_ff99bsc0*χ*_OL3_*eζ*_OL1_ forcefield, as it consistently showed a larger mismatch with experimental data. Our goal was to adjust the free-energy difference between C3’-endo and C2’-endo conformations for methylated uridine from the original +0.5 kcal/mol to -0.2 kcal/mol, in line with experimental findings.

This correction was implemented by applying an additional grid bias to the relevant dihedral coordinate. The bias was carefully designed to selectively tune the populations of each conformational basin while minimally perturbing the free-energy barrier between them, thus preserving the exchange kinetics across the two conformers (see Methods).

The additional potential energy term (see bias in Figure 3) successfully corrects the imbalance between the two ribose conformations, restoring the experimentally observed populations for the isolated methylated uridine. In the absence of experimental data for purines, we assumed that this correction could be applied to the A-1 puckering angle in the methylated ribozyme. All other forcefield parameters were left unchanged.

**Figure 3:**
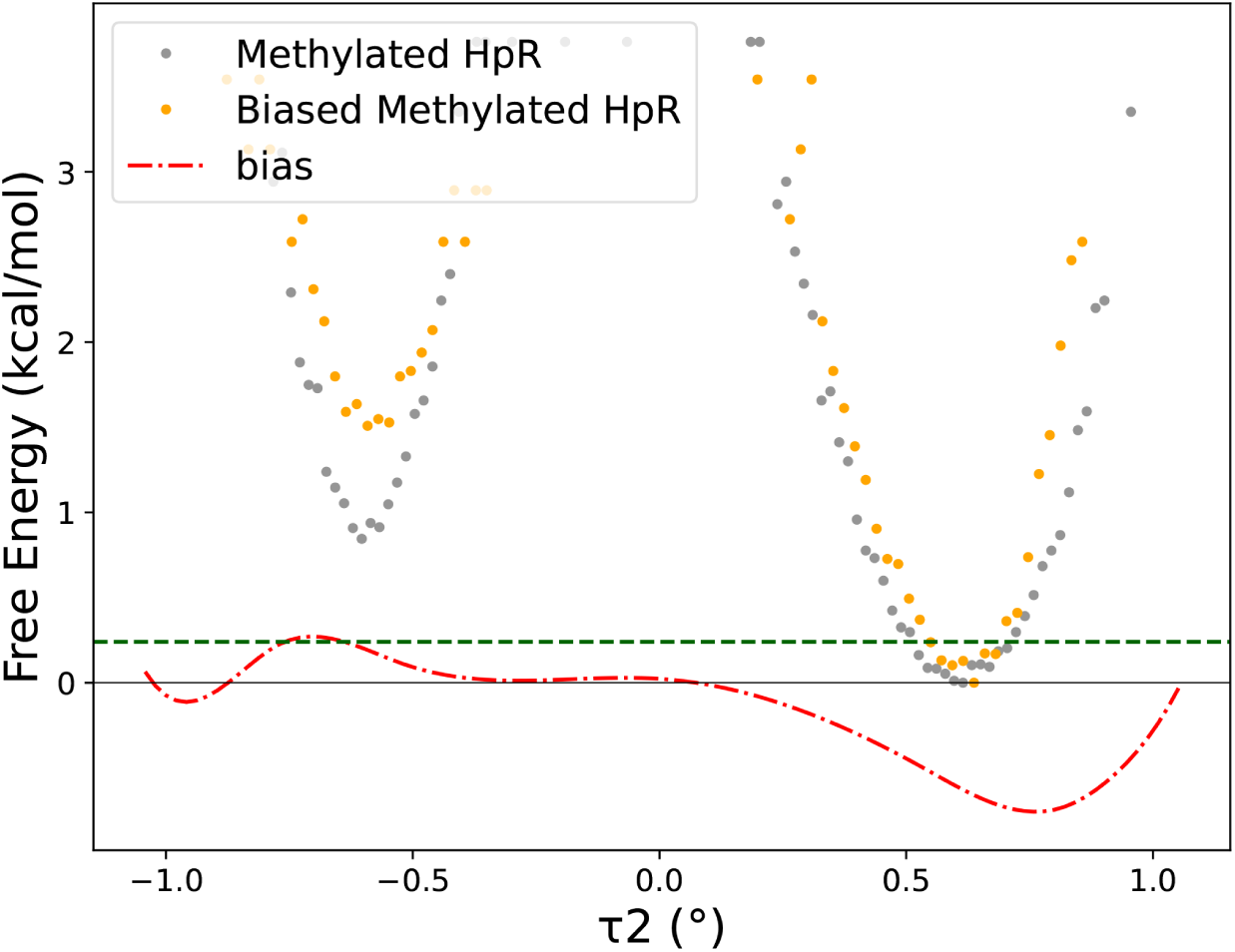
Potentials of mean force along *τ*_2_ dihedral angles in the two REST2 simulations of the HpR structure with 2’-OMe modification: with the original Amber14_ff99bsc0*χ*_OL3_*eζ*_OL1_ parametrization, with (in orange) and without (in grey) the restraint applied on A-1 *τ*_2_. C2’-endo conformations correspond to negative values of the *τ*_2_ angle, and C3’-endo to positive values. See Methods for more details on how this restraint is built.

As shown in Figure 3, the corrected forcefield shifts the free-energy landscape along the A-1 *τ*_2_ dihedral angle in the methylated ribozyme. Consequently, the population ratio between the C3’-endo and C2’-endo conformations increases, and the free-energy difference between the two conformers as well (Table 3). As expected, even after correcting the forcefield, methylation continues to strongly stabilize the C2’-endo conformation relative to the unmethylated case (Table 1). This indicates that correcting for potential forcefield biases does not alter the core conclusion of our study.

**Table 3:** *p_C_*_3_*t_−endo_/p_C_*_2_*t_−endo_* ratios and the corresponding Δ*G* values calculated from the *τ*_2_ dihedral distribution in the converged regions of various HpR REST2 simulations,comparing different variants with forcefield modifications.The standard deviation was evaluated from time block analysis, with 50 ns blocs.

| System | Parametrization | C3'-endo/C2'-endo | $\Delta G$(kcal/mol) |
| --- | --- | --- | --- |
| Solution demethylated PC | $\chi_{OL3}\epsilon\zeta_{OL1}$ with add. dih. term | inf<sup>1</sup> | < -4<sup>1</sup> |
| Solution methylated PC | $\chi_{OL3}\epsilon\zeta_{OL1}$ + local correction | 21 ± 13 | - 1.7 ± 0.4 |

Error bars for this run could not be obtained as in the C2'-endo conformation was never observed in the trajectory, resulting in infinite ratios.

### 3.5 Reparametrization of all pseudo-rotation angles

Having examined the limitations of current RNA forcefields in accurately capturing the effects of methylation on ribose puckering, and explored possible corrective strategies, we now shift focus to the unmethylated case. This scenario is ultimately the most stringent test, as it reflects the ribozyme’s pre-catalytic state.

When comparing simulation results for the uridine RNA mononucleoside with experimental data, we observed subtle but systematic deviations in the free-energy difference between the C2’-endo and C3’-endo conformations. The DES-Amber 3.20 forcefield closely matched experimental values, whereas Amber14_ff99bsc0*χ*_OL3_*eζ*_OL1_ deviated by 0.3 kcal/mol and, more critically, incorrectly favored the C2’-endo conformation.

To address this discrepancy, we introduced a systematic correction to the ribose puckering terms by adding an extra dihedral potential energy term, analogous to the earlier grid bias but now incorporated directly into the forcefield. Our aim was to assess whether this adjustment would meaningfully impact both the conformational landscape and the active site geometry of the ribozyme. Full details of the parametrization are provided in the Methods section, and the specific parameters used are listed in the Supporting Information.

As shown in Figure S8, this reparameterization does not significantly affect the overall eRMSD of the ribozyme structures. An analysis of the RMSF for individual residues confirmed that local dynamics remained largely unchanged, with only minor fluctuations observed upon forcefield modification. While these results warrant more extensive investigation, they suggest that the correction does not degrade the integrity of the original forcefield.

Notably, the C2’-endo conformation of the A-1 ribose is no longer observed in our REST2 simulations using the corrected forcefield. This implies that the free-energy difference between the two puckering states now likely exceeds that of the unmodified Amber14_ff99bsc0*χ*_OL3_*eζ*_OL1_ (Table 3). Once again, we emphasize that conformational preferences are influenced not only by the intrinsic enthalpic tendencies encoded in the forcefield (and aligned here with experimental benchmarks), but also by environmental context and broader forcefield features, such as electrostatics and neighboring dihedral terms. In particular, the DES-Amber 3.20 forcefield, which required no correction due to its accurate puckering balance when not methylated, yields a 2 kcal/mol difference relative to Amber14_ff99bsc0*χ*_OL3_*eζ*_OL1_.

### 3.6 Molecular origins for the stabilization of A-1 puckering conformations

We now demonstrate that our simulations provide molecular-level insights into the stabilization of the C2’-endo conformation upon methylation. To explore this, we clustered the ribozyme conformations sampled during the unperturbed REST2 simulations using the YACARE algorithm ^80^ (see Methods), applied to the Amber14_ff99bsc0*χ*_OL3_*eζ*_OL1_ forcefield data. This allowed us to identify representative structures for each puckering state (Figure 4).

**Figure 4:**
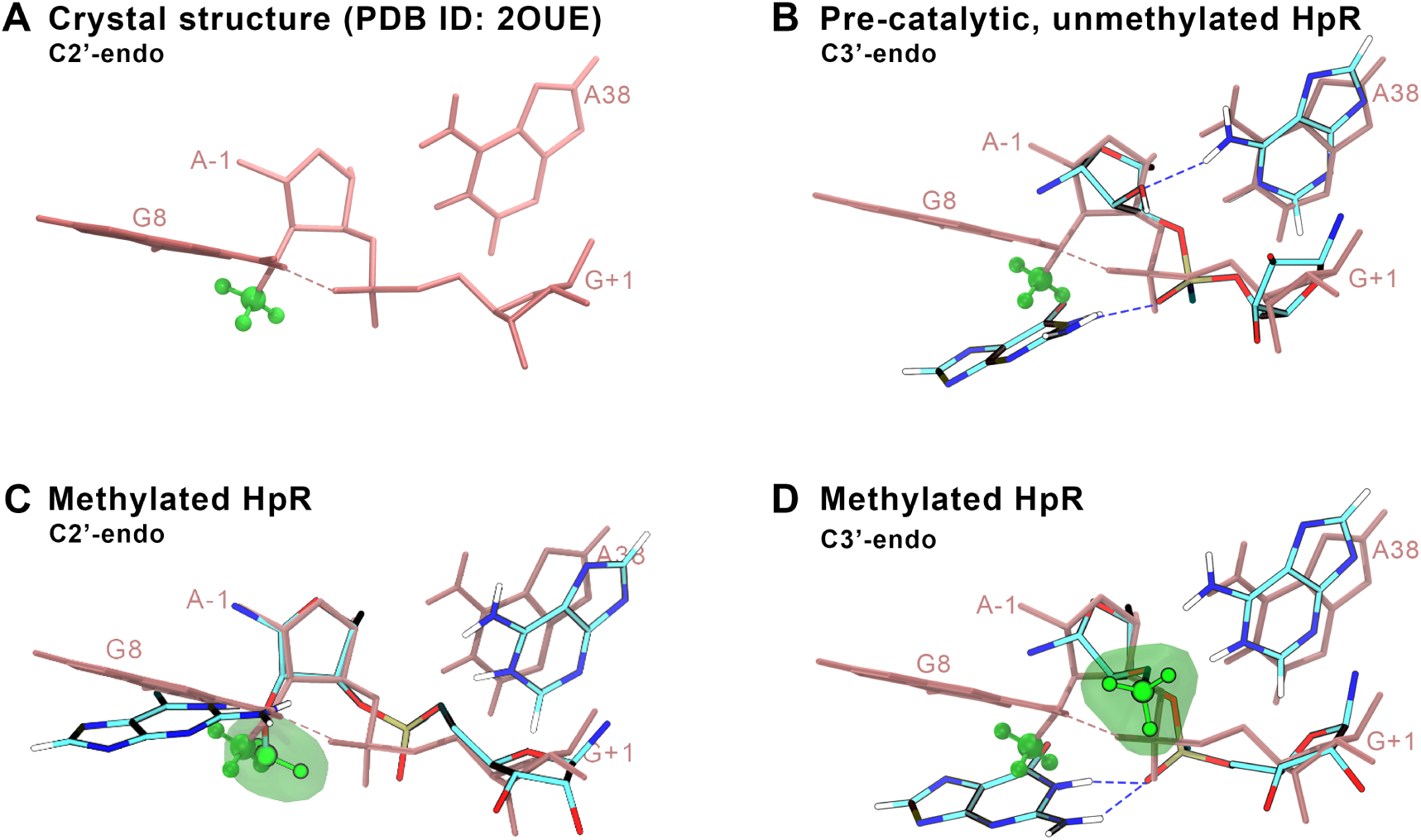
Active site structure of the HpR in its most representative conformations, with the scissile junction formed by G+1 and A-1 (their nucleobases are not represented) and the surrounding A38 and G8 nucleobases. A) Crystallographic structure which was used as base structure for all systems (PDB ID : 2OUE). This structure, here on its own, is superposed in pink in B,C,D to help distinguishing the divergence of the different sampled conformats. (B, C, D, E) Structural organization of the main C2’-endo (C) and C3’-endo (B,D) conformations extracted from our clustering analysis of the HpR simulations, methylated (C,D) and unmethylated (B). The methyl group is highlighted in green, with its volume visible in transparent green. The key H-bonds involving the nucleophilic O2’ and the non-bridging oxygens of the junction appear as dashed blue lines.

We compared these representative conformations to the crystallographic (methylated) structure (Figure 4A). In the absence of the methyl group, the A-1 ribose consistently adopts a C3’-endo conformation, which induces significant rearrangements within the active site (Figure 4B). These include notable changes in the ribose geometry, phosphate linkage orientation, and the positioning of the catalytically important G8 residue. In this configuration, multiple hydrogen bonds (H-bonds) form: the hydroxyl group at A-1:O2’ acts both as a proton donor—interacting with non-bridging phosphate oxygens—and as a proton acceptor—interacting with the nearby A38 nucleobase. This flexibility enables G8 to shift closer to the scissile junction, establishing an H-bond with one of the phosphate’s non-bridging oxygens.

Upon methylation, the C2’-endo conformation observed in the crystal structure becomes stabilized relative to the hydroxylated (unmethylated) case (Figure 4C). The representative active-site structure in this conformation more closely resembles the crystallographic state, with the exception of local rotations of the scissile phosphate. Notably, both G8 and A38 maintain positions similar to those in the crystal structure, and the methyl group itself remains relatively stable throughout the simulation. By contrast, in the methylated C3’-endo conformation, the active site adopts a geometry similar to that of the hydroxylated form, except that A-1:O2’ can no longer participate in hydrogen bonding (Figure 4D).

Based on these findings, we propose that the stabilization of the C3’-endo conformation in the hydroxylated system may be driven, at least in part, by favorable hydrogen bonding involving A-1:O2’ and key catalytic residues such as G8 and A38. In the methylated system, the absence of these interactions, combined with possible steric hindrance between the methyl group and A38, may instead favor the C2’-endo conformation. In this latter state, the methyl group positions itself beneath the ribose plane (Figure 4A and C), while important hydrogen bonds between G8 and the scissile phosphate are preserved.

### 3.7 Effect of methylation in other ribozymes

We now assess whether our results are specific to the HpR or if they can be generalized to other ribozymes. Structural data from various ribozymes indicate that the methylated reactive sugar often adopts a C2’-endo conformation ^62,86,111–113^, a feature considered as maybe important for understanding ribozyme reactivity ^111^. However, in all available cases, experimental structures were typically obtained using either deshydroxylation or methylation, modifications that, as discussed earlier, can significantly alter the ribose puckering equilibrium.

To test generalizability, we selected two additional small self-cleaving ribozymes, each with distinct characteristics, that we simulated with Amber14_ff99bsc0*χ*_OL3_*eζ*_OL1_. The first, the hammerhead ribozyme (HhR), shares several structural and mechanistic features with the HpR and is well-studied while remaining computationally accessible. The second, the glmS ribozyme, requires a sugar activator (glucosamine-6-phosphate) for self-cleavage and is considerably larger than HpR and HhR. Its overall fold and active-site architecture differ significantly. In both systems, crystallographic structures typically feature methylation at the reactive 2’-OH, and the corresponding ribose adopts a C2’-endo conformation ^62,86^.

Starting from the methylated crystal structure of the HhR (PDB 3zp8) ^86^, we carried out long, single-replica unbiased MD simulations in both the methylated and unmethylated states at the reactive C17 sugar. Although magnesium is required for catalysis, we excluded it from our simulations because (i) it was not resolved in the crystal structure, and its precise placement is nontrivial, and (ii) it is thought to act primarily on the leaving group side (O5’ of C1.1), with limited expected influence on the puckering of the C17 ribose. Remarkably, in the methylated system, the C2’-endo conformation remained stable over 500 ns. In contrast, the unmethylated ribose rapidly and irreversibly flipped to a C3’-endo conformation, mirroring the behavior observed in the HpR (Figure S2A and B).

To further enhance conformational sampling, we performed 500-ns REST2 simulations on the methylated HhR (Figure S2C). These simulations revealed a gradual equilibration between the C2’-endo and C3’-endo states, with final populations closely matching those observed in the HpR (Δ*G* = Δ1.4 kcal/mol). This similarity is consistent with the local structural context: although the hammerhead active site is more solvent-exposed than that of the HpR, it shares key features on the nucleophilic side. In particular, G12 in the HpR occupies a position analogous to G8 in the HpR (Figure S9). Consequently, in the unmethylated state, the C3’-endo conformation is stabilized by a network of hydrogen bonds involving G12, the non-bridging oxygens of the scissile phosphate, O2’, and A11 – this last nucleobase reminding the interactions of A38 for the HpR. In contrast, the C2’-endo conformation (and the methylated C3’-endo state) disrupts this interaction network.

Methylation also significantly affects the ribose conformation in the glmS ribozyme, although via a distinct mechanism. Due to the system’s large size, REST2 simulations were impractical, so we conducted 800-ns unbiased MD simulations in both the methylated and unmethylated states. These were initialized from the methylated crystal structure (PDB 3g8s) in the C2’-endo conformation ^62^. In contrast to HhR and HpR, the cleavage site in glmS is located at the terminus of an RNA strand (Figure S10), and catalysis requires a sugar cofactor, which was not present in the crystal structure used a as starting point for our simulations (PDB 3g8s) as we focused on the apo, pre-catalytic state.

In the methylated form, the C2’-endo conformation remained stable throughout the trajectory (Figure S11A). However, in the absence of the methyl group, the ribose shifted within tens of nanoseconds to a C4’-exo/O4’-endo conformation (Figure S11B). The active site architecture in glmS differs radically from those of the other ribozymes, involving two key residues (G33 and G57) that form hydrogen bonds with the 2’-OH. These interactions may explain the ribose’s atypical puckering behavior (Figure S10). In the methylated state, the C2’-endo conformation likely minimizes steric clashes within the active site, contributing to its stability.

## 4 Discussion

O2’ methylations are found in several key RNA systems. For many naturally-occuring RNAs, methylation of the 2’OH group is a common post-translational modification (PTM) that regulates RNA function ^22–25,25–39^. In addition, structural studies of self-splicing ribozymes often require the inactivation of the ribose O2’ group, which serves as an intramolecular nucleophile for self-splicing ^10,11^. While the literature generally suggests that methylation reinforces the preference of RNA ribose for the C3’endo puckering conformation ^40,50–54^, our findings, based on molecular dynamics (MD) simulations with enhanced conformational space exploration, provide evidence that this trend can be reversed. Specifically, we show that a single-site methylation in the reactive sugar of the hairpin ribozyme can preferentially stabilize the C2’-endo conformation relative to the C3’-endo sugar puckering, while the unmodified nucleotide (which cannot be directly studied experimentally) exhibits a relative preference for the C3’-endo conformation. We show that this trend is robust against change in the forcefields, and greatly exceed comparatively small inaccuracies of the forcefields to quantitatively describe the puckering equilibrium. Upon fine tuning the forcefield to reproduce experimental data on isolated mononucleosides using two different strategies, we show that this trend remains valid. Our estimations of the C3’-endo/C2’-endo free-energy difference evolution upon methylation for these different conditions and systems are recapitulated in a schematic diagram (Figure 5).

**Figure 5:**
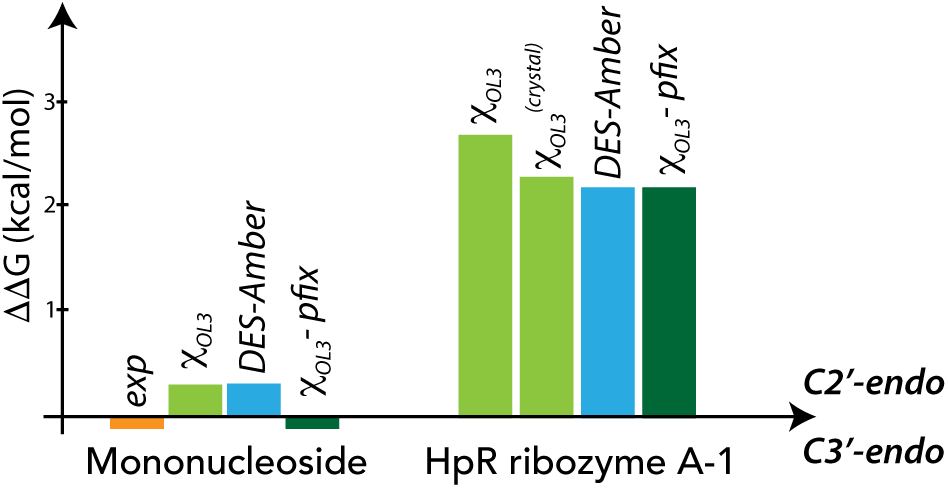
Free-energy change upon methylation for the isolated mononucleoside (left) and the HpR ribozyme A-1 sugar (right). *χ*_OL3_ refers to Amber14_ff99bsc0*χ*_OL3_*eζ*_OL1_ forcefield, *χ*_OL3_ − *pfix* to the grid bias used to reproduce experimental data on isolated 2’-O-methylated uridine, and *χ^crystal^* to the crystal simulation (all other systems are solvated in an aqueous environment).

Our results may seem at odds with previous experimental data ^40,50–54^. However, we argue that the claim that methylation reinforces the stability of C3’endo conformations has been overstated. First, the effect is, in fact, subtle from a freeenergy perspective. Earlier reports suggesting a stabilization of 0.1-0.8 kcal/mol are based on enthalpic considerations alone, which are largely compensated by entropic effects ^40,50^. We also note that quantum calculations, which reproduce this trend, tend to overestimate the effects, reporting (enthalpic) C3’-endo stabilizations of 2-5 kcal/mol relative to C2’-endo ^53,54^. This discrepancy likely arises from limited sampling in these calculations along collective variables (CVs) orthogonal to the puckering angle.

That being said, it is clear that methylation can reinforce the stability of duplex regions in RNA structures, particularly for residues already engaged in base-pairing or those at the edge of base-paired regions ^40,59^. Methylation may also stabilize the formation of duplexes from initially unpaired residues. However, C2’-endo conformations are not uncommon for methylated ribose. For example, ribozymes often exhibit C2’-endo conformations for the methylated reactive sugar, which has been regarded as crucial for understanding their reactivity ^111^. Recent high-resolution structures of the human ribosome provide a more nuanced view of the conformational shift induced by methylation ^69–71^. For example, in a structure with 59 methylated ribose ^69^, 9% of the methylated residues adopt a C2’-endo conformation.

These considerations suggest that the occurence of C2’-endo conformations in a methylated residue is not surprising, and could be context-dependent (O2’ methylation stabilizes duplex regions in C3’-endo conformation but may at the same time favor local arrangements that are more stable in a C2’-endo conformation). However, what is perhaps more unexpected is that, upon removal of the methyl group, there is a relative stabilization of the C3’endo conformation by approximately 2-2.5 kcal/mol. We demonstrate that this effect is robust across changes in the forcefield, and that for two other ribozymes, hammerhead and glmS, methylation also has profound structural impacts in the active site. Furthermore, although we observed systematic deviations from the experimental reference data regarding the puckering angle, local and systematic corrections to the forcefield, calibrated on isolated uracil nucleotides, did not lead to any change in this behavior. Taken together, while we cannot rule out complex forcefield artifacts, these results provide solid support for the observed behavior.

Interestingly, while in situ experimental observation of the puckering equilibrium in conditions where the ribozyme is active remains out of reach unless a new method for controlling the self-cleaving reaction is developed, it is worth noting that several experimental structures have achieved inactivation of the 2’OH group by dehydroxylation ^11,16,17^ or by mutating other active site residues, most of which are thought to be crucial for the reaction ^11,18,21^. In all of these structures, the puckering angle is not in the C2’-endo conformation.

Our results have important implications. They demonstrate that the effect of chemical modifications in the active site, particularly on the reactive ribose, may induce structural changes that have often been overlooked, although some of the distortions induced by methylation had been already suggested ^65^. This is especially surprising since such modifications can also be regulatory, and are considered crucial in many other contexts. It is expected, however, since local RNA structures extensively rely on hydrogen bond patterns, which are inevitably affected by these modifications.

This highlights that structural interpretations of the mechanism should be carefully considered, as the methylated structure may not a good proxy of the precatalytic state. If the C3’-endo conformation is indeed the most favored conformation of the A-1 ribose, it could provide additional insights into the reaction mechanism, as recently discussed ^68^. Specifically, this geometry would be more favorable for a phosphate-assisted reaction pathway, such as the monoanionic mechanism, which is actually the mechanism for the uncatalyzed reaction ^114,115^. In contrast, the hydrogen bond pattern required for a general acid/general base mechanism appears unlikely. Further exploration of this discrepancy could offer deeper understanding into the precise catalytic mechanism of the ribozyme.

## 5 Conclusions

In this work, we use molecular dynamics simulations to provide microscopic evidence of the changes induced by 2’-O-methylation in the reactive ribose of the precatalytic state of the hairpin ribozyme, and demonstrate similar effects for the hammerhead and glmS ribozymes. Methylation is a commonly used strategy to chemically impair reactivity for structural studies, but our simulations suggest that it induces a local conformational change in the active site that critically affects the sugar puckering in connection with the hydrogen-bond network and local steric hindrance.

Our results challenge and refine the common assumption that methylation enhances the stability of the C3’-endo conformation relative to the C2’-endo conformation in comparison to unmethylated RNA. We show that methylation can, in fact, have the opposite effect depending on the local nucleic acid environment. We also argue that the magnitude of this effect has often been overstated in the literature, as it has been clearly demonstrated primarily for duplex motifs. For isolated, non-interacting mononucleotides, the effect is extremely mild from a free-energy perspective, not exceeding 0.2 kcal/mol. Moreover, recent high-resolution structures of the human ribosome contain dozens of methylated residues, a fraction of which is in the C2’-endo conformation ^69–71^.

While forcefield inaccuracies may improperly balance the populations in each conformational basin, our findings are consistent across two different forcefield combinations. Additionally, we propose both specific and systematic reparametriza-tions of the ribose dihedral term in the Amber14_ff99bsc0*χ*_OL3_*eζ*_OL1_ forcefield, calibrated using isolated pyrimidine nucleotide data, which leads to the same quantitative conclusions. This demonstrates that the stabilization of one conformation over another cannot be understood purely from the dihedral energy landscape; rather, it is likely governed by interactions with the local environment.

Our work also calls for caution when interpreting the precatalytic structure of ribozymes obtained through chemical modifications. The observed alteration of the sugar puckering may have important consequences for the mechanism of the catalyzed chemical reaction. In this regard, the crystal structure may be misleading, and careful relaxation through atomistic molecular dynamics simulations combined with enhanced sampling techniques should precede any subsequent reactivity studies.

Finally, we note that *in situ* measurements of the active site structure are experimentally challenging due to the self-reactive nature of the biomolecule. Time-resolved spectroscopy techniques may provide indirect information about the local conformations of the active site during the catalytic step, offering further validation of the effects of methylation. In the meantime, this work demonstrates that atomistic molecular dynamics simulations can provide experimentally inaccessible details that are crucial for understanding ribozyme reactivity, while remaining in agreement with sparse and indirect data from other systems.

## 7 Author contribution statement

S.F. and G.S. conceived the research. S.F. conducted the simulations, S.F. and G.S. analysed the results. S.F. and G.S. wrote and reviewed the manuscript.

## 8. Acknowledgments

We thank Élise Duboué-Dijon for discussions. The research leading to these results has received funding from the European Research Council under the European Union’s Eighth Framework Program (H2020/2014-2020)/ERC Grant Agreement No. 757111 (G.S.). The simulations presented here benefited from access to the HPC resources of TGCC under the allocation A0130811005 made by GENCI (Grand Equipement National de Calcul Intensif).

## 9 Data availability statement

All simulation inputs and data used to generate figures will be made available on Zenodo. Analysis scripts will be made available in the GitHub page of this paper upon final acceptance of this paper. A tutorial with some related protocols is available from the PLUMED-tutorials website ^116^. All simulations were performed with standard open source software as described in the methods.

## Supporting information

### Simulation parameters

The OL force fields can be found online at: https://fch.upol.cz/ffol/index.php. The DESAmber 3.20 force field parameters in Gromacs format were obtained by contacting, modified so that the energy reference for dihedral terms was consistent with that typically used in the original forcefield. This version of the DES-Amber 3.20 force field can be found in the zenodo public repository of our previous study (https://zenodo.org/records/11033971).

### Non-standard nucleotides parameters

We give below the atomic charges for the MRA residue (methylated adenosine), which were obtained by adding to the DES-Amber adenine residue the charge difference between RA and MRA residues in other Amber14_ff99bsc0 force fields. Doing so, we generate the modified adenine residue by adding the charge redistribution suggested in previous studies. Please note that this has not been properly recalculated nor optimized within the framework of this force field.

### Modified MRA residue parameters used in trajectories parameterized with DES-Amber 3.20 (rtp format)

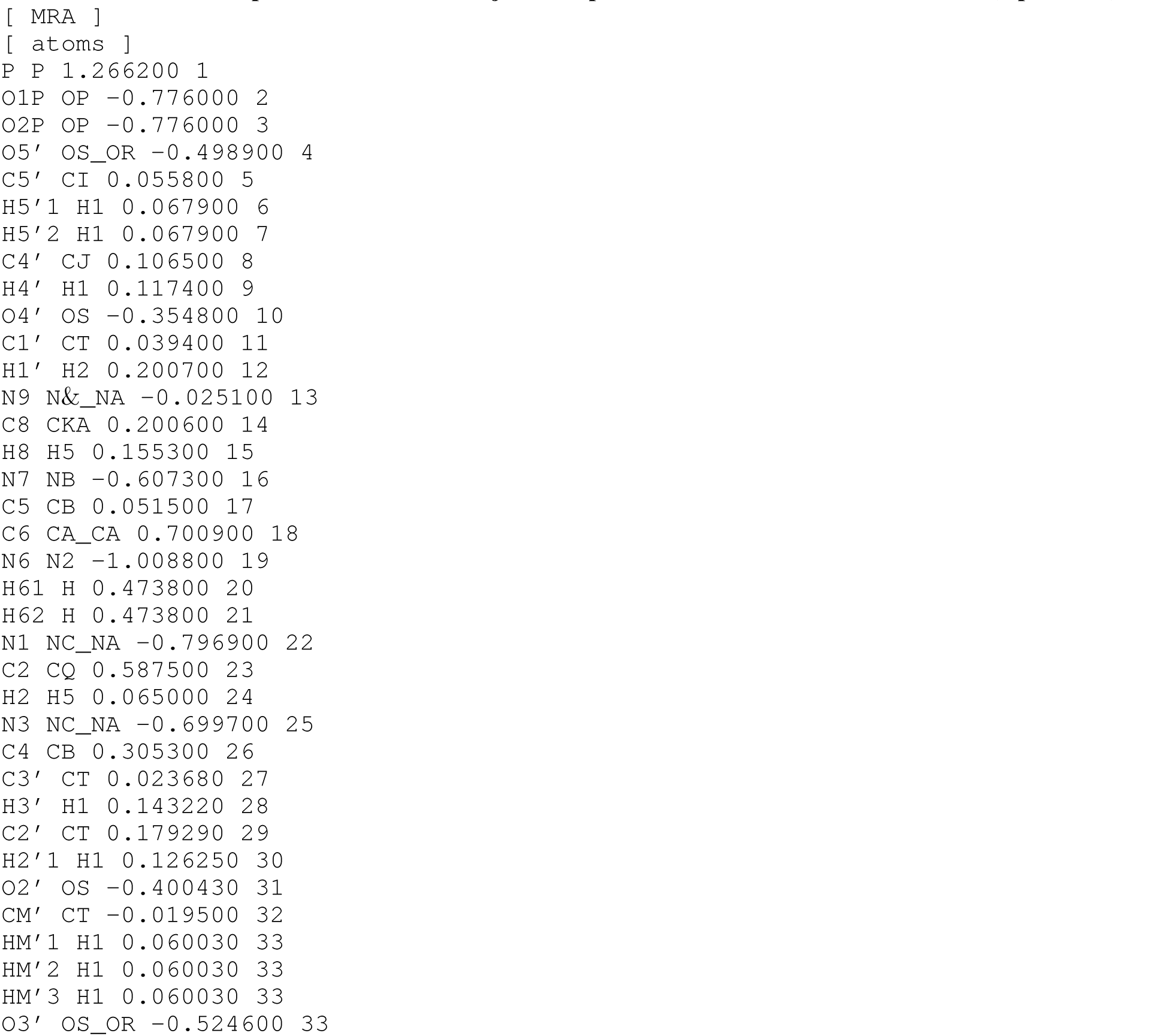

### Simulating Mononucleotides and Dinucleotides

**Modified MUN residue parameters used in trajectories parameterized with Amber14_ff99bsc0***χ*_OL3_*eζ*_OL1_ **(rtp format)**.

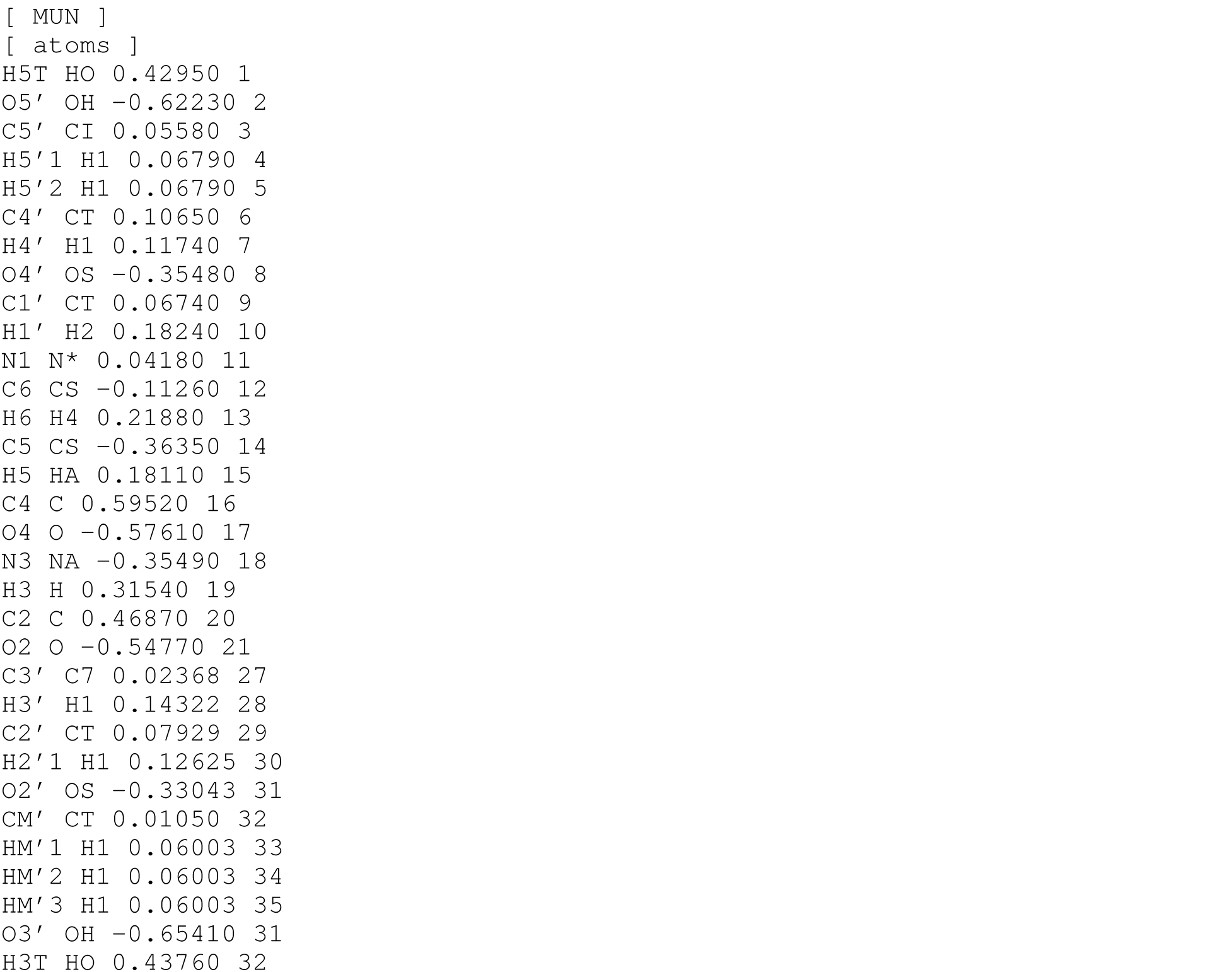

### Modified MUN residue parameters used in trajectories parameterized with DES-Amber 3.20 (rtp format)

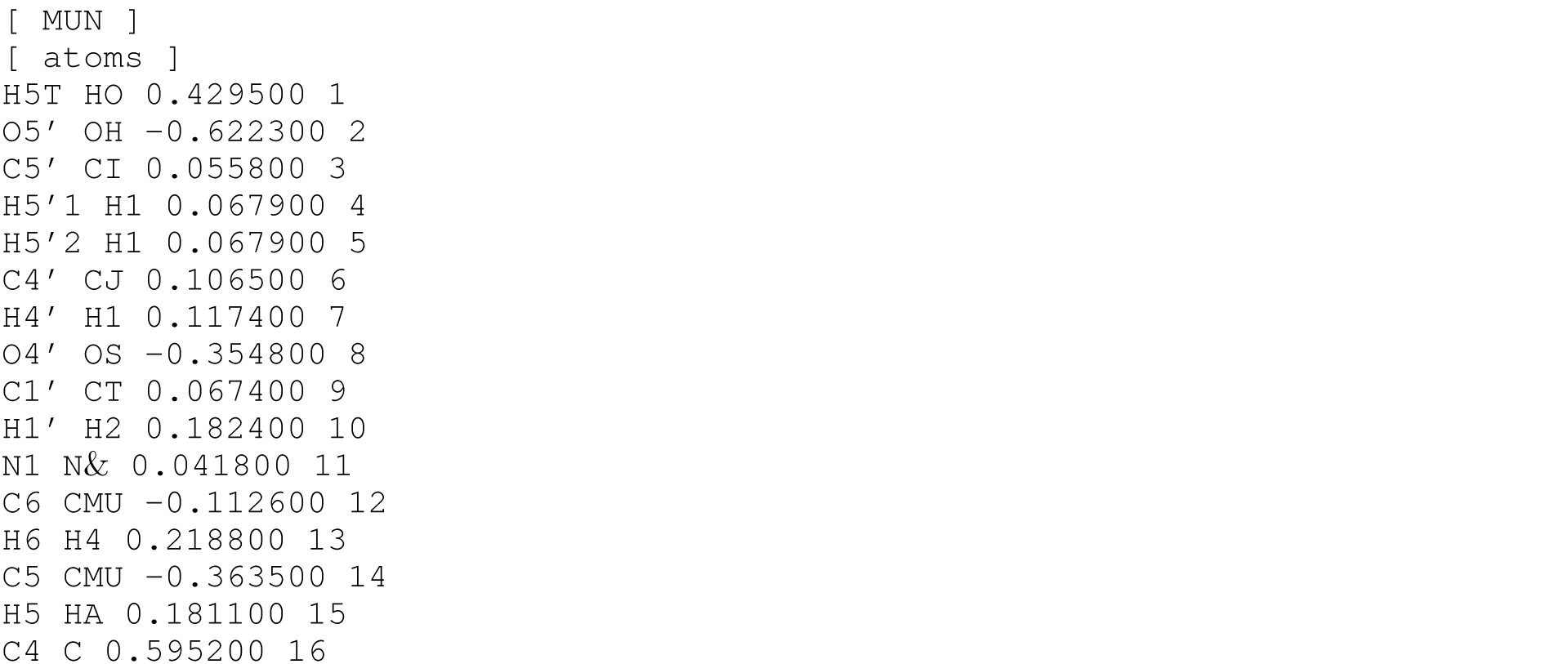
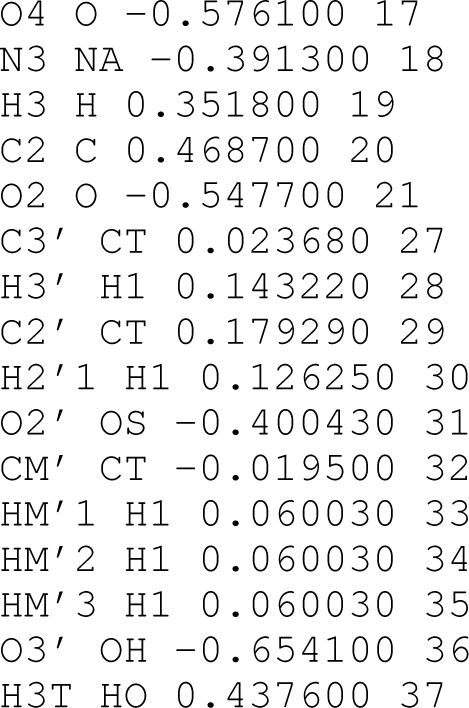

### Puckering Rescaling

The following parameters are added on top of the existing dihedral terms in Amber14_ff99bsc0*χ*_OL3_*eζ*_OL1_. The profile of the added terms can be seen in Figure S6.

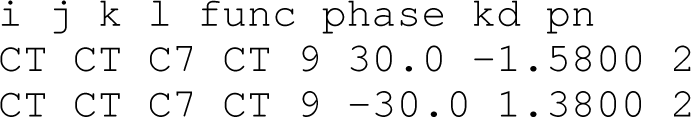

---

**Figure S1:**
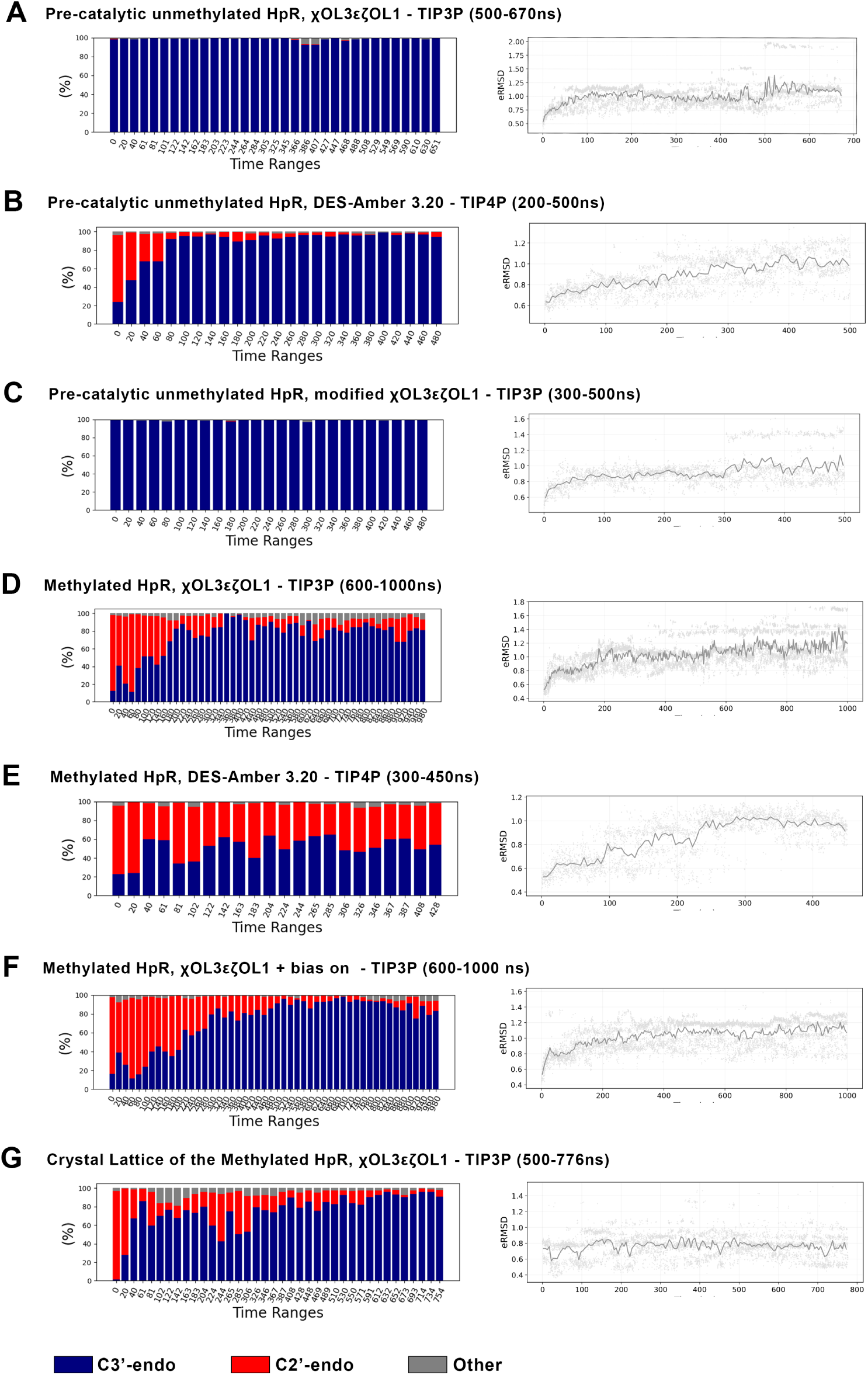
Local and global observables of the unscaled replica used to determine convergence for the Hairpin Ribozyme REST2 simulations of this study. The left panel shows the evolution of the proportion of the puckering pseudo-rotation angle C2’-endo and C3’-endo populations along time in the lowest replica trajectory. The right panel shows the evolution in time of the structure eRMSD ^93^ relative to each respective initial structure. We indicate within parentheses the trajectory portion taken for analysis.

**Figure S2:**
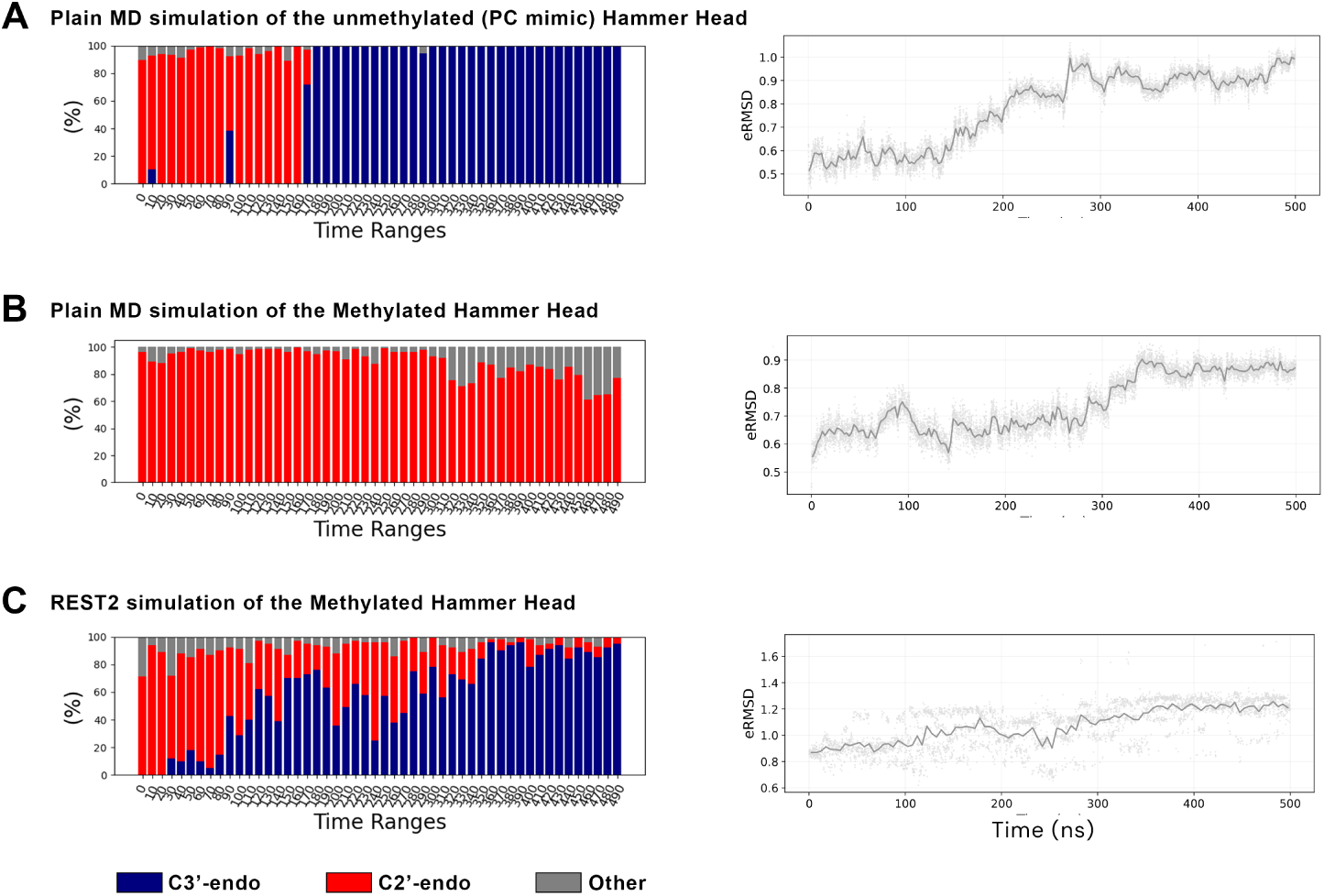
Local and global observables of the unscaled replica used to determine convergence for the Hammerhead REST2 and plain MD simulations of this study. The left panel shows the evolution of the proportion of the puckering pseudo-rotation angle C2’-endo and C3’-endo populations along time in the lowest replica trajectory. The right panel shows the evolution in time of the structure eRMSD ^93^ relative to each respective initial structure.

**Figure S3:**
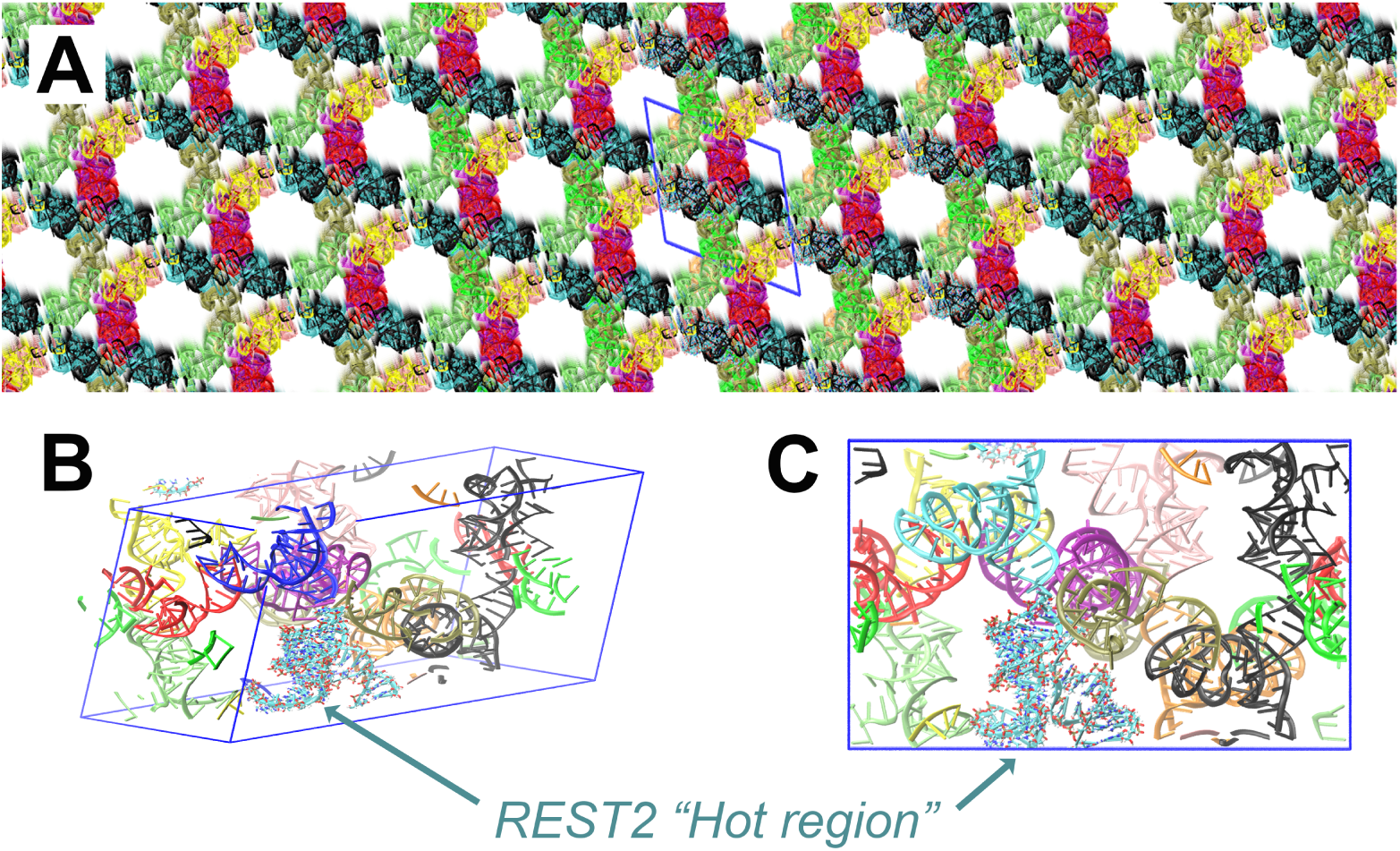
Several views of the crystal lattice of the hairpin ribozyme, constructed from the PDB entry 2OUE structure. (A) Side view of the crystal lattice and its periodic images to highlights the symmetry of the lattice. The triclinic simulation box appears in blue, and the colors match those in (B) and (C). (B, C) Two views of the triclinic simulation box, with the twelve hairpin ribozymes constituting one unit cell shown in different colors. The “hot region,” represented by one of the twelve HpRs, is shown in cyan and displayed in full-atom “licorice” style (a feature of VMD ^78^). All other eleven ribozymes are treated as solvent molecules in the REST2 framework.

**Figure S4:**
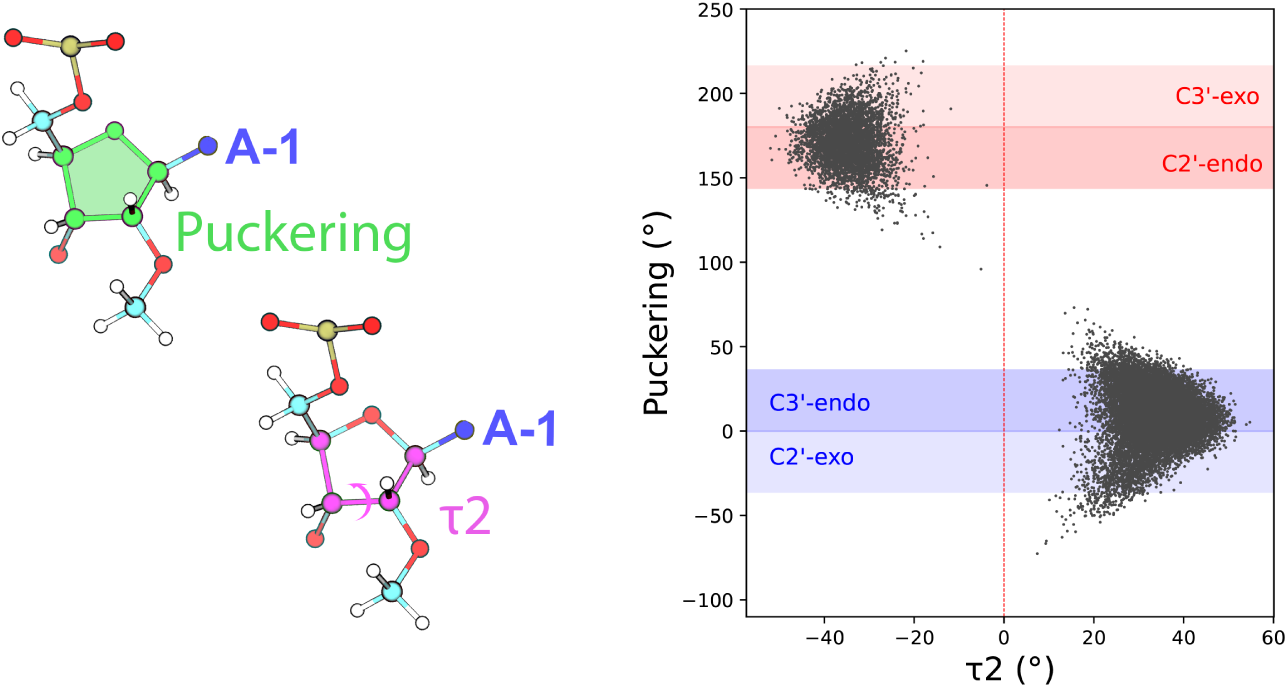
Correlation between *τ*_2_ and the puckering pseudo-rotation angle of A-1 observed in the REST2 simulation of the methylated hairpin ribozyme in the bulk (Amber14_ff99bsc0*χ*_OL3_*eζ*_OL1_ forcefield), over a -110° to +250° range. *τ*_2_ ≤ 0 corresponds to the “southern” puckering conformation (C2’-endo and C3’-exo), while *τ*_2_ ≥ 0 aligns with the “northern” puckering conformation (C3’-endo and C2’-exo).

**Figure S5:**
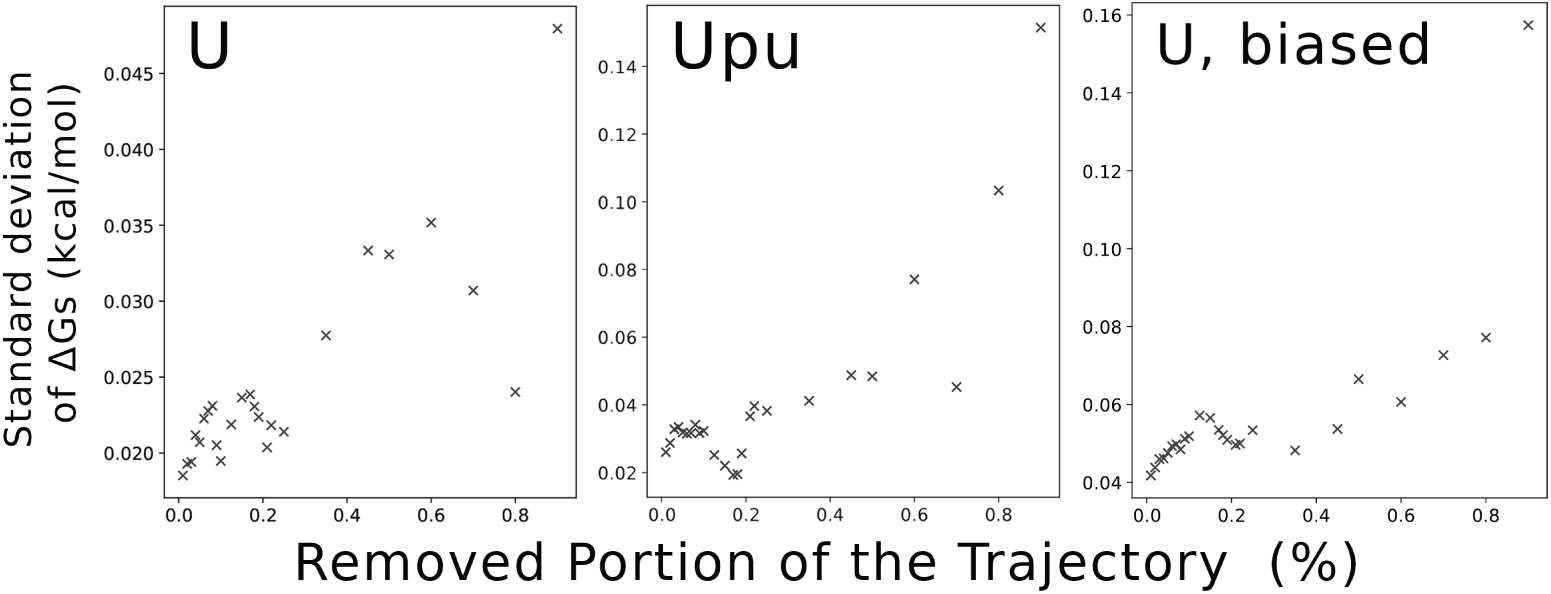
The *p_C_*_3_*t_−endo_/p_C_*_2_*t_−endo_* ratio standard deviation calculated over truncated 5-replica OPES trajectories of different sizes, for U, biased U, and UpU. The x-axis represents the percentage of the trajectory removed. This analysis was performed to evaluate the to-be-removed equilibration time of the OPES simulations.

**Figure S6:**
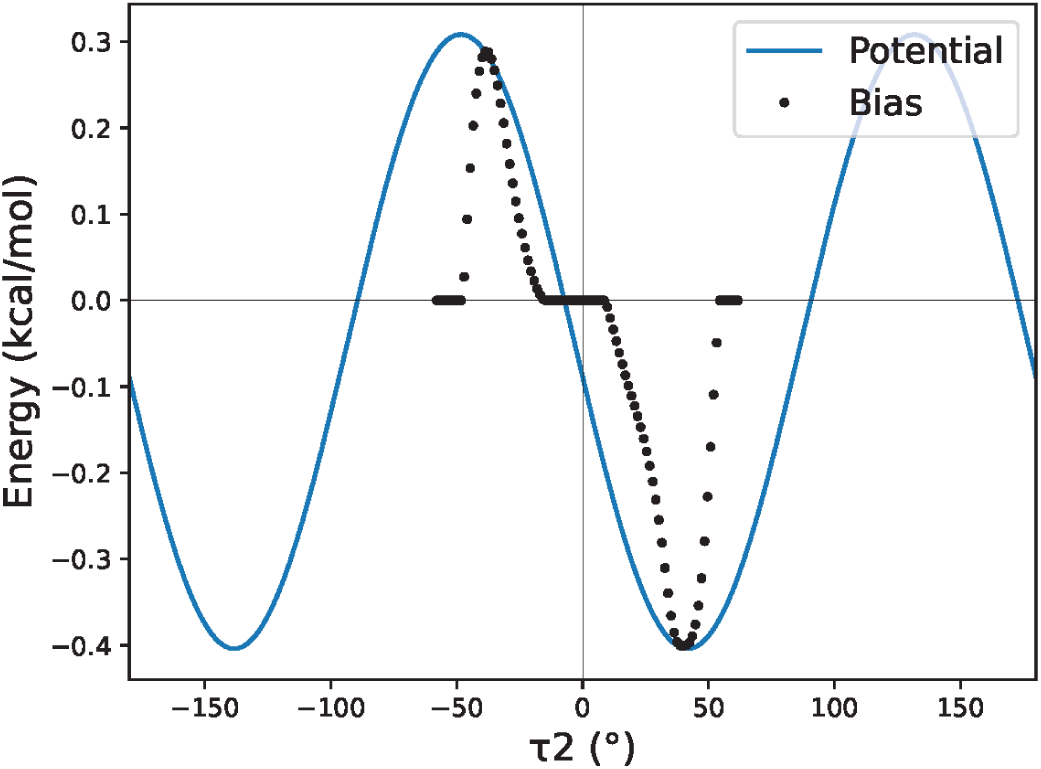
Potential energy profile of the two additional dihedral terms which will be applied to all riboses (in blue), together with the bias grid (in black dots) they were built from. More details regarding the construction of the bias grid can be found in the Methods section.

**Figure S7:**
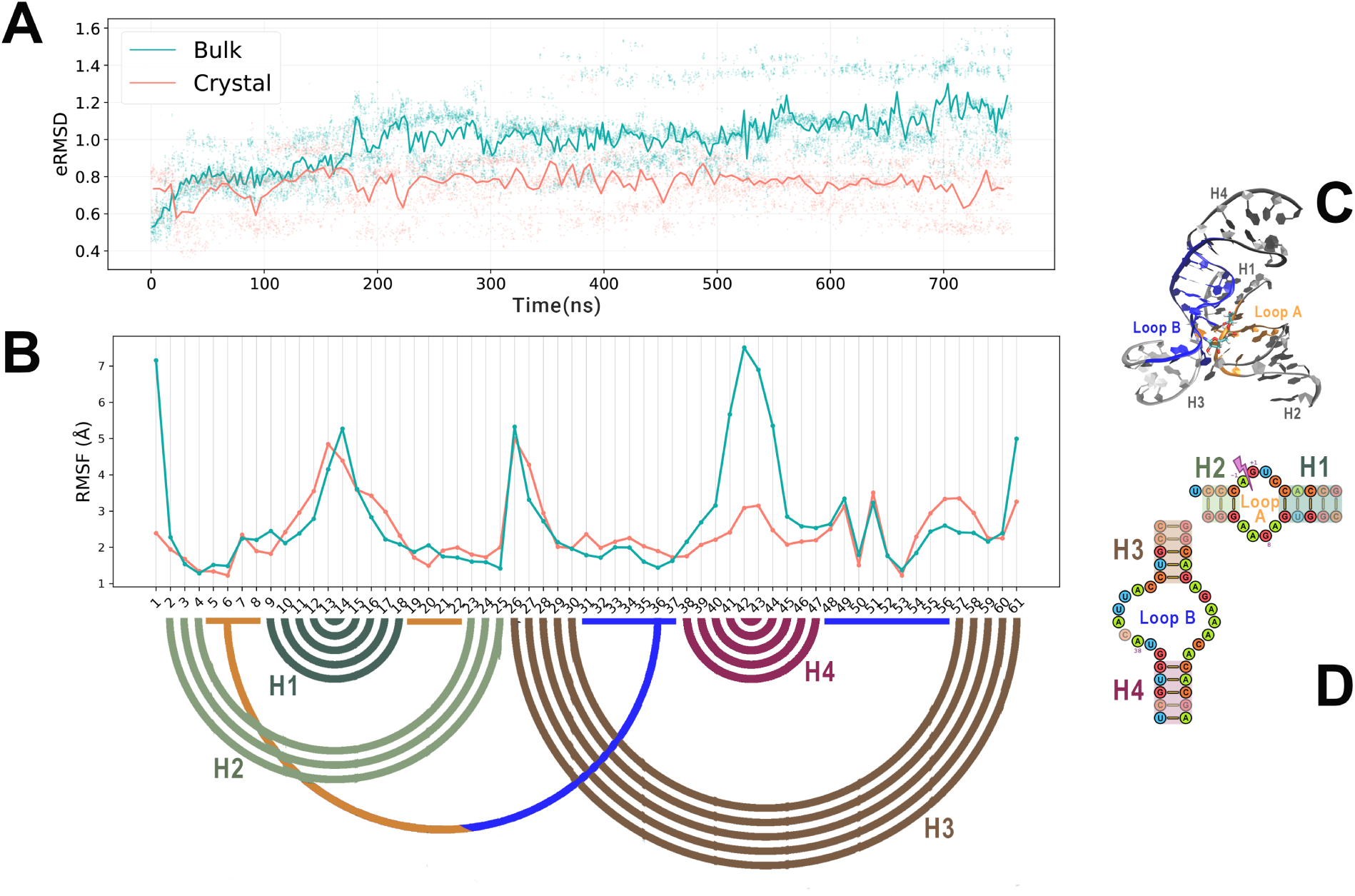
Compared stability of the methylated hairpin ribozyme in crystal (in orange) versus bulk (in turquoise) environment (Amber14_ff99bsc0*χ*_OL3_*eζ*_OL1_ forcefield). (A) eRMSD time evolution, calculated from the native structure (PDB entry 2OUE). From this plot, convergence times where estimated at 200 ns for both simulations. (B) Per residue RMSF calculated on the C1’ atoms and on the last 500 ns of each simulation, together with a bow scheme representing the native Watson and Crick base pairs, colored accordingly to (C), the 3D view of the native structure, and (D), its secondary structure.

**Figure S8:**
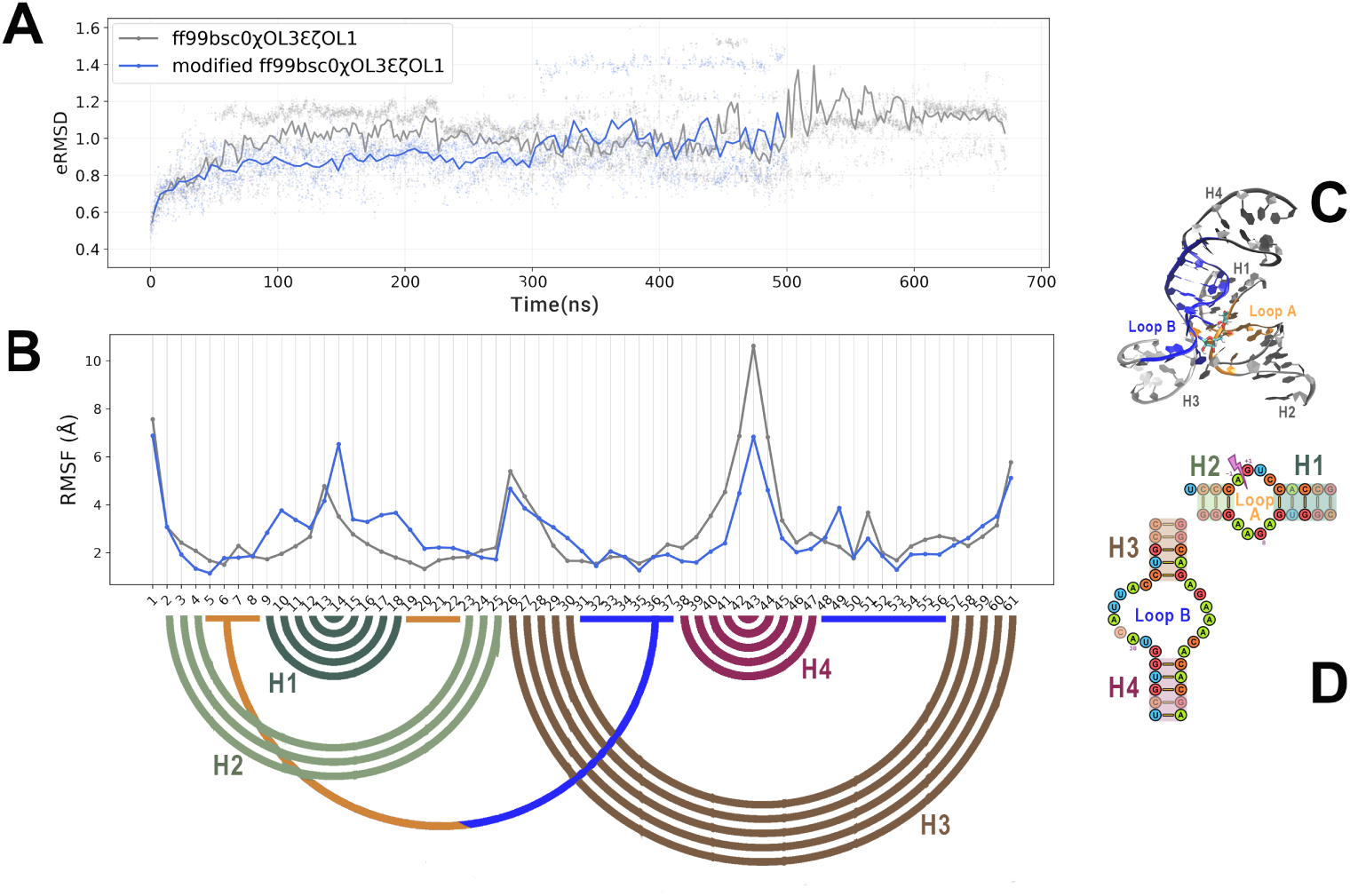
Compared behavior of the pre-catalytic state of the hairpin ribozyme when parametrized with the original Amber14_ff99bsc0*χ*_OL3_*eζ*_OL1_ (in gray) and the same force field with the additional dihedral terms (in blue). See the methods section for more details on the construction of the additional dihedral terms. (A) eRMSD time evolution, calculated from the native structure (PDB entry 2OUE). From this plot, convergence times where estimated at ∼ 500 ns for the original ff and ∼ 300 ns for the corrected ff simulation. (B) Per residue RMSF calculated on the C1’ atoms and on the converged part of each simulation, together with a bow scheme representing the native Watson and Crick base pairs, colored accordingly to (C), the 3D view of the native structure, and (D), its secondary structure.

**Figure S9:**
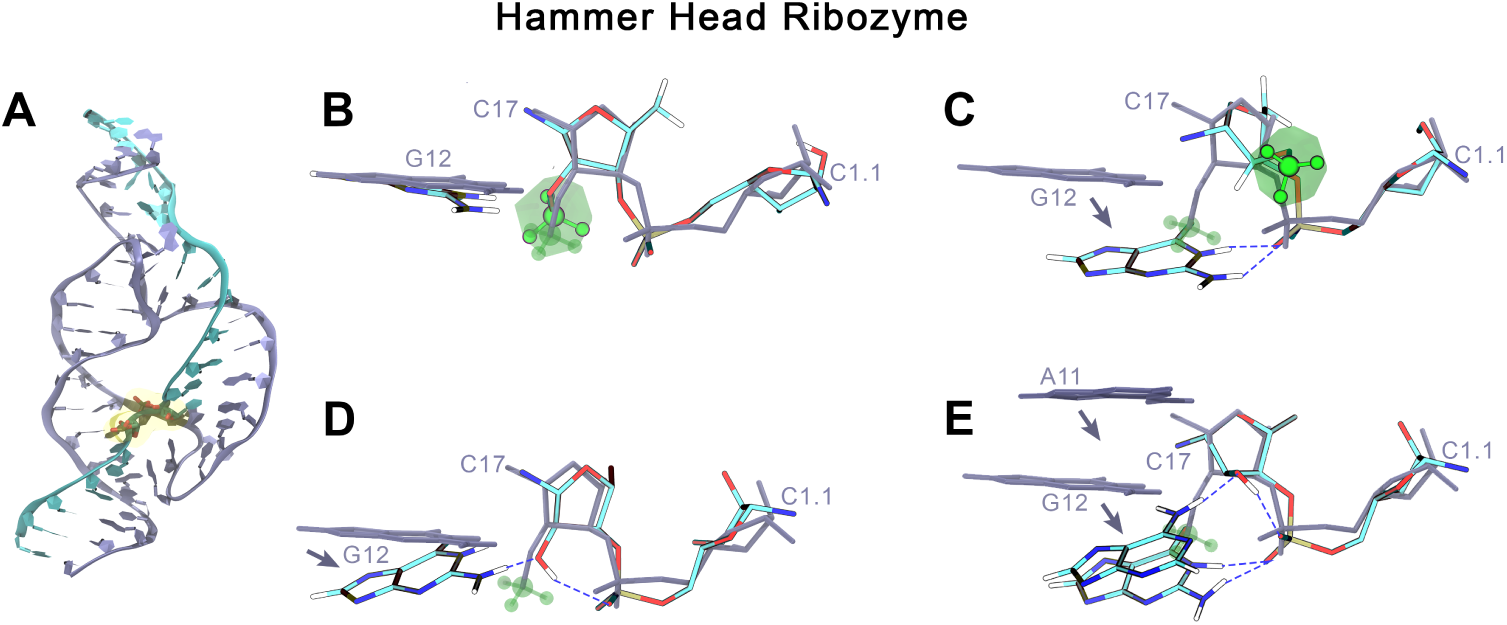
A) A 3D view of the entire Hammerhead ribozyme simulated in this study, with the catalytic site highlighted in yellow and its atoms fully represented in licorice VMD representation. The RNA strand upon which cleavage occurs is in blue. (B, C, D, E) Structural organization of the main C2’-endo (B, D) and C3’-endo (C, E) conformations extracted from our clustering analysis of the Hammerhead simulations, methylated (B, C) and unmethylated (D, E). The PDB structure from which we built the initial conformations is superimposed in blue in each view to help distinguish the diverging features of the local configurations of the catalytic site. The methyl group is highlighted in green, with its volume visible in transparent green. The key H-bonds involving the nucleophilic O2’ and the non-bridging oxygens of the junction appear as dashed blue lines.

**Figure S10:**
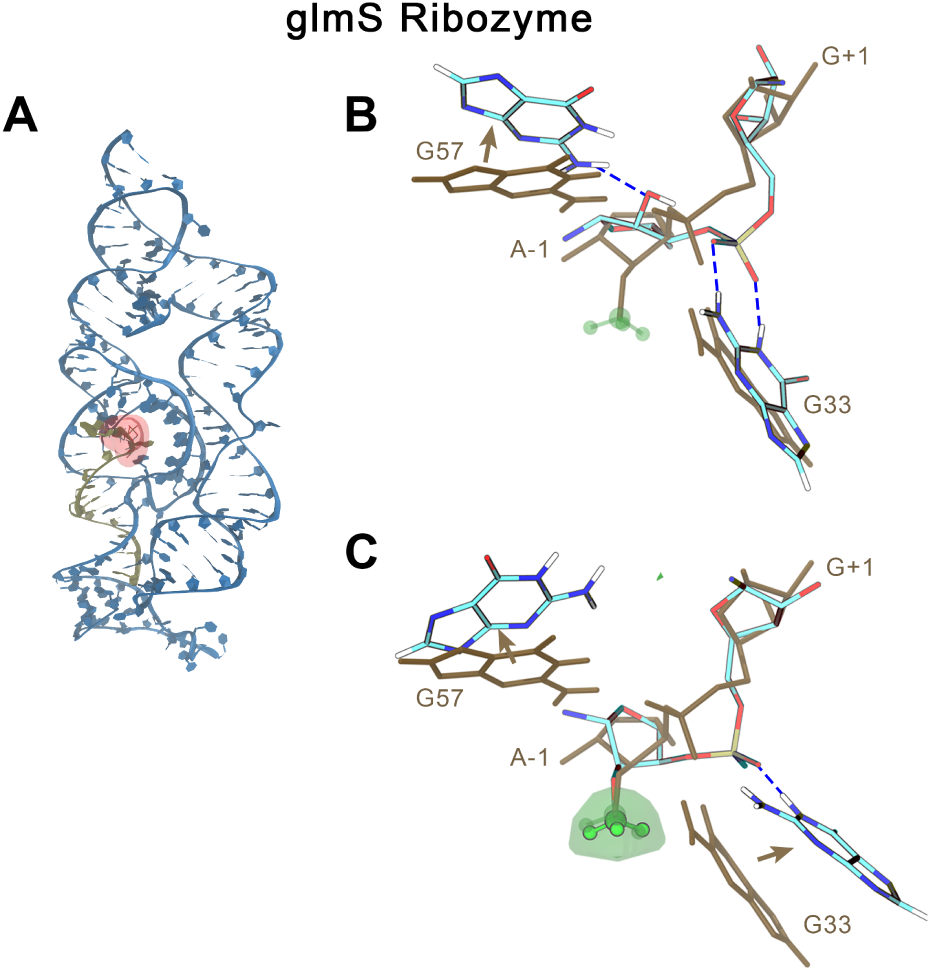
(A) A 3D view of the entire glmS simulated in this study, with the catalytic site highlighted in red and its atoms fully represented in licorice VMD representation. The RNA strand upon which cleavage occurs is in yellow. (B, C) Structural organization of the main conformations extracted from our clustering analysis of the glmS simulations, unmethylated (B) and methylated (C). The PDB structure from which we built the initial conformations is superimposed in brown in each view to help distinguish the diverging features of the local configurations of the catalytic site. The methyl group is highlighted in green, with its volume visible in transparent green. The key H-bonds involving the nucleophilic O2’ and the non-bridging oxygens of the junction appear as dashed blue lines.

**Figure S11:**
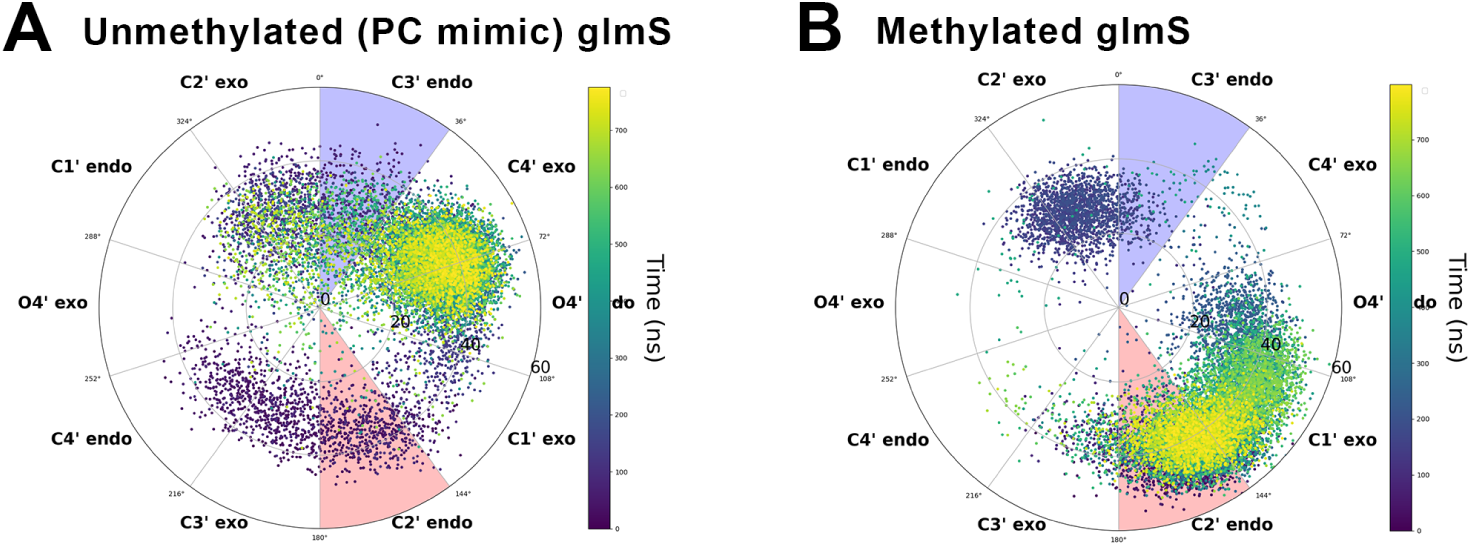
Puckering conformation distribution of the plain MD of the methylated (A) and unmethyated (B) glmS ribozyme simulations. The distribution is time-colored.

